# Hidden Talents: Silent Gene Clusters Encoding Magnetic Organelle Biosynthesis in a Non-Magnetotactic Phototrophic Bacterium

**DOI:** 10.1101/2022.04.19.488322

**Authors:** M.V. Dziuba, A. Paulus, L. Schramm, R.P. Awal, M. Pósfai, C.L. Monteil, S. Fouteau, R. Uebe, D. Schüler

**Affiliations:** Department of Microbiology, Faculty of Biology, Chemistry and Geosciences, University of Bayreuth, Germany; Department of Microbial Biochemistry, Faculty of Life Sciences: Food, Nutrition and Health, University of Bayreuth, Germany; Research Institute of Biomolecular and Chemical Engineering, University of Pannonia, Veszprém, Hungary; Aix-Marseille University, CEA, CNRS, Biosciences and Biotechnologies Institute of Aix-Marseille, Saint Paul lez Durance, France; LABGeM, Genomique Metabolique, CEA, Genoscope, Institut Francois Jacob, CNRS, Universite d’Evry, Universite Paris-Saclay, Evry, France

**Author notes:** The authors to whom correspondence should be addressed (D. Schüler), (R. Uebe).

## Abstract

Horizontal gene transfer is a powerful source of innovations in prokaryotes that can affect almost any cellular system, including microbial organelles. However, typically rapid loss of non-functional gene acquisitions obscures the mechanisms determining the fate of horizontally transferable genes. Here, we report the first discovery of a horizontally inherited gene cluster encoding biosynthesis of magnetosomes, the organelles used by magnetotactic bacteria for navigation, in a non-magnetotactic phototrophic bacterium *Rhodovastum atsumiense*. We show that these clusters were inactivated through transcriptional silencing and antisense RNA regulation, but retain functionality, as several genes were able to complement the orthologous deletions in a remotely related magnetotactic bacterium. The laboratory transfer of foreign magnetosome genes to *R. atsumiense* was found to endow the strain with magnetosome biosynthesis, but strong negative selection led to rapid loss of this trait upon subcultivation. Our results provide insight into the horizontal dissemination of gene clusters encoding complex prokaryotic organelles and illuminate the potential mechanisms of their genomic preservation in a dormant state.

## Introduction

The discovery of various prokaryotic organelles during the last decades has led to the rejection of the previously common view that bacterial cells have simplistic organization^1^. One of the best-studied examples of such organelles are magnetosomes synthesized by magnetotactic bacteria (MTB). Magnetosomes consist of magnetic iron oxide or sulphide cores enveloped by a bilipid layer and are aligned in one or multiple chains within the cells^2^. They enable MTB to passively align with the Earth’s magnetic field lines, which in combination with active cellular movement and a highly complex signal transductory aerotactic network facilitates their search for optimal redox conditions (magnetotaxis) at the oxic-anoxic transition zones of the stratified aquatic habitats, which MTB ubiquitously populate^3,4^. The ability to exert precise biological control over the synthesis of these highly ordered mineral-containing structures has placed MTB in the spotlight of a growing number of studies focused on the genetics, molecular mechanisms, evolution, and biotechnological application of bacterial magnetic biomineralization^2,5–7^.

Formation of a membranous compartment, magnetic crystal synthesis and assembly of magnetosome chains require the coordinated action of >30 magnetosome-associated proteins, which are encoded within several magnetosome gene clusters (MGCs). The evolutionary history of MGCs still represents a conundrum. Intriguingly, the ability to form magnetosomes is widely scattered on the bacterial tree of life: MTB are affiliated with *Alpha-, Delta-, Gamma-, Lambda-, Zeta*- and *Etaproteobacteria* classes of phylum *Proteobacteria*, as well as in the phyla *Nitrospirae, Nitrospinae, Omnitrophica, Latescibacteria, Planctomycetes, Fibrobacteres*, and *Riflebacteria*^8,9^. Based on the broad phylogenetic distribution of MTB and the diversity of magnetosome crystal composition and morphology, multiple evolutionary origins of magnetotaxis had initially been proposed^10^. However, recent data indicate that at least the core set of magnetosome genes emerged only once and is highly conserved among all MTB^8,11,12^. Nevertheless, as to when they emerged and what was the further course of their evolution, is still a matter of vigorous debates^13,14^. On the one hand, phylogenetic trees built from 16S rRNA or phylogenetic marker proteins are largely congruent with those based on the shared magnetosome proteins, suggesting potential vertical inheritance from a common ancestor before the emergence of the major phyla^12^. However, this scenario places the origin of this highly complex trait very close to the last common bacterial ancestor (LBCA), questioning the view on ancient prokaryotes as organisms with the primitive cellular organization. Additionally, this scenario would imply numerous losses of magnetotaxis genes during phylogenetic divergence, which is not a parsimonious explanation, and assume only marginal importance to the role that HGT could play in the MGCs evolution^13,14^. On the other hand, multiple violations of the phylogenetic tree congruency were observed, indicating instances of horizontal gene transfer (HGT)^9,15–19^. Moreover, the potential ability for MGCs mobilization and transfer is supported by the fact that in many of the described MTB, MGCs are found in genomic regions of plasticity, so-called genomic magnetosome islands (MAI), which is supported by the deviating G + C content, codon adaptation index (CAI), tetranucleotide frequency and abundance of mobile elements^20,21^. Furthermore, the ability of some hitherto non-magnetotactic organisms to biomineralize magnetosomes after receiving MGCs was recently documented applying synthetic biology methods. Thus, the laboratory transfer of the major MGCs from a model MTB *Magnetospirillum gryphiswaldense* MSR-1 to a purple photosynthetic alphaproteobacterium *Rhodospirillum rubrum* and a closely related but non-magnetotactic *Magnetospirillum* sp. indeed endowed both species with magnetosome biosynthesis^22,23^. These experiments indicated the sufficiency of the transferred gene set for magnetosome formation and the overall ability of these foreign hosts to integrate the magnetosome biosynthesis pathway into their metabolic networks, suggesting that this might also occur under natural conditions.

However, to which extent HGT impacts the evolution of magnetosomes is not yet clear. It is generally assumed that the horizontally transferred genes remain fixed in the population if they are properly expressed and confer advantages to the host, i.e., improve their fitness^24^. Hence, the currently documented MTB diversity supposedly represents the product of natural selection for fixation of magnetosome formation as a favourable trait, whereas all the non-functional or non-beneficial gene acquisitions have been eliminated from the gene pool and therefore have remained unavailable for analysis or laboratory experiments to date.

Here, we report the discovery of the silent, yet functional MGCs in the culturable non-magnetotactic phototrophic bacterium *Rhodovastum atsumiense* G2-11, which were apparently acquired by a recent HGT from an alphaproteobacterial MTB. We further present a comprehensive study of the transcriptional pattern, functionality, and fitness effect of the magnetosome genes in G2-11. Moreover, we ‘magnetized’ G2-11 through artificial HGT of the major MGCs from MSR-1 under laboratory conditions and explored the trait stability upon subcultivations. Our results provide the first evidence for horizontal dissemination of gene clusters encoding bacterial magnetic organelles outside MTB and illuminate the potential mechanisms of their preservation in a latent state.

## Results

### The phototrophic species Rhodovastum atsumiense G2-11 acquired MGCs from an unknown alphaproteobacterial MTB by recent HGT

In a systematic database search for novel MGCs, we identified several orthologs of known magnetosome genes in the recently released draft genome sequence of the culturable anoxygenic phototroph *Rhodovastum atsumiense* G2-11^25^. This finding was unexpected as, after isolation of G2-11 from a paddy field more than 20 years ago, no magnetosome formation has been reported^26^. Furthermore, no MTB has been identified so far among phototrophs or within the *Acetobacteraceae* family to which G2-11 belongs^26^ (Fig. 1a).

**Fig. 1.**
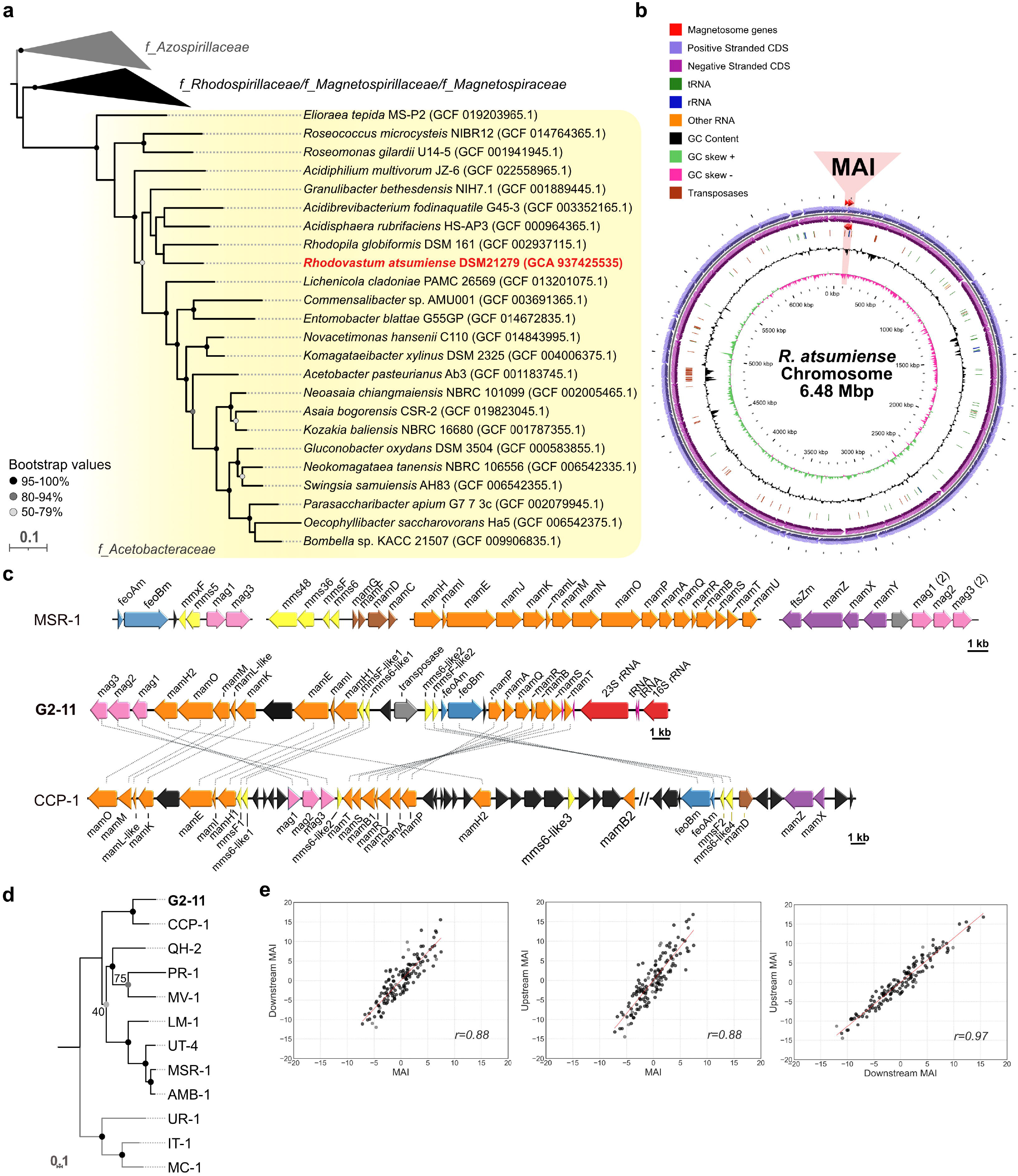
Phylogeny, chromosome and MGCs organization of G2-11. (a) The maximum likelihood phylogenetic tree based on ribosomal proteins demonstrates position of G2-11 (highlighted in red) within family *Acetobacteraceae*. The *Azospirillaceae* family was used as an outgroup based on the latest *Alphaproteobacteria* phylogeny. Branch length represents the number of base substitutions per site. Values at nodes indicate branch support calculated from 500 replicates using non-parametric bootstrap analysis. Bootstrap values <50% are not shown. The families are labelled according to GTDB taxonomy. (b) Circular map of the G2-11 chromosome. The region of magnetosome genomic island (MAI) is highlighted in red. (c) Organization of the G2-11 MGCs in comparison to MSR-1 and the uncultivated MTB CCP-1. Magnetosome genes are colored according to their position within the magnetosome operons of MSR-1. Connecting dotted lines indicate synteny between the G2-11 genes and CCP-1. (d) Maximum likelihood phylogenetic tree of concatenated amino acid sequences of the shared MamKMOPAQBST proteins for G2-11 and the MTB strains: uncultivated calcium carbonate producing MTB CCP-1, *Magnetospira* sp. QH-2, *Ca*. Terasakiella magnetica PR-1, *Magnetovibrio blakemorei* MV-1, *Ca*. Magneticavibrio boulderlitore LM-1, *Magnetospirillum* sp. UT-4, *M. gryphiswaldense* MSR-1, *M. magneticum* AMB-1. *Ca*. Magnetaquicoccus inordinatus UR-1, *Magnetococcus marinus* MC-1, and *Magnetofaba australis* IT-1 were used as an outgroup. Branch length represents the number of base substitutions per site. Values at nodes indicate branch support calculated from 500 replicates using non-parametric bootstrap analysis. (e) Distribution of z-normalized tetranucleotide frequencies in the G2-11 MAI region in comparison to the flanking regions (Upstream MAI and Downstream MAI). Each dot represents a tetranucleotide combination. Pearson’s correlation coefficients (r) are shown on the graphs.

Since the available genome version (Accession Number NZ_VWPK00000000) was highly fragmented (226 contigs) and the magnetosome genes were distributed over several contigs with gaps, we first re-sequenced and assembled the complete genome using long reads generated by Illumina HiSeq and Nanopore technologies. The resulting genome consists of one chromosome (6.48 Mb) and eight plasmids ranging from 10,690 bp to 220,129 bp in size (Fig. 1b, Supplementary Figure S1). The putative magnetosome genes localize within a single region (27.5 kb) on the chromosome, compactly organized in four operon-like clusters comprising the following genes: *mag123*, (*mms6-like1*)(*mmsF-like1*)*mamH1IEKLMOH2*, (*mms6*-like2)(*mmsF*-like2), and *feoAmBm-mamPAQRBST* (Fig. 1c). They include all genes thought to be essential for magnetosome biosynthesis (*mamIELMOQB*)^27^ and appear to be intact, as no obvious frameshifts or nonsense mutations could be detected. Protein alignments using BLASTP suggested that the closest orthologues of the magnetosome genes from G2-11 are found among alphaproteobacterial MTB (Supplementary Table S1). The phylogenetic analysis of concatenated amino acid sequences of the magnetosome proteins MamKMOPAQBST demonstrated that the sequences from G2-11 reliably cluster with those of an uncultivated calcium carbonate producing MTB CCP-1^28^ (Fig. 1d). The low completeness of the CCP-1 genome does not allow us to reliably infer the relationship between G2-11 and CCP-1 using phylogenomic markers, e.g., ribosomal proteins, as we conducted for the *Acetobacteraceae* family (Fig. 1a). Nonetheless, CCP-1 has been shown to belong to *Azospirillaceae* based on the 16S rRNA tree^28^, which occupies an outgroup position related to *Rhodospirillaceae* and *Acetobacteraceae*, according to the latest *Alphaproteobacteria* phylogeny^29^. This incongruence in the phylogenetic positions of these two species and their magnetosome genes is best explained by the HGT of the MGCs in G2-11 from an unknown MTB belonging to *Azospirillaceae*.

Although the magnetosome genes from G2-11 and CCP-1 have a close phylogenetic relationship, the comparative analysis of their MGCs revealed considerable differences in the organization. First, G2-11 lacks several accessory magnetosome genes (*mamX, mamZ* and *mamD*), which were previously shown to be universally present in alphaproteobacterial MTB and, although being not essential, are important for proper magnetic crystal formation in MSR-1 and *M. magneticum* AMB-1^30,31^. Their absence in G2-11 could be explained by functional differences in the magnetosome biosynthesis pathways, incomplete horizontal transfer of the MGCs, or a secondary loss of these genes in G2-11. Furthermore, the MGCs of CCP-1 are interspersed by >20 genes with no homology to known magnetosome genes (Fig. 1c). In contrast, the compact MGCs in G2-11 contain only a few genes that could not be associated with magnetosome biosynthesis.

Tetranucleotide usage patterns are frequently employed as a complementary tool to group organisms since they bear a reliable phylogenetic signal^32^. Likewise, deviations of tetranucleotide usage in a certain fragment from the flanking genome regions can indicate HGT^21^. Comparison of the z-normalized tetranucleotide frequencies of the MGCs (27.5 kb) with the flanking upstream (117.7 kb) and downstream (79.5 kb) fragments showed a considerably lower correlation between them (Pearson’s correlation coefficient r=0.88 with both flanking fragments) than between the flanking fragments themselves (Pearson’s r=0.97, Fig. 1e). This indicates a significant difference in the tetranucleotide composition of the MGCs compared to the flanking genomic regions and supports a foreign origin of the magnetosome genes in G2-11 suggested by the phylogenetic analysis. Besides, the presence of a mobile element (transposase) and position of the MGCs directly downstream of a tRNA gene, a common hotspot for integration of genomic islands^33–35^, suggests that the MGCs of G2-11 are indeed located on a genomic island, the so-called magnetosome island (MAI), like in many other MTB^20,21^. Unfortunately, the lack of other representatives of the genus *Rhodovastum* makes it impossible to infer whether the MAI was transferred directly to G2-11 or the last common ancestor of the genus. However, its compact organization and conspicuous tetranucleotide usage suggest a relatively recent HGT event.

### G2-11 does not form magnetosomes under laboratory conditions

Although magnetosome genes discovered in G2-11 comply with the minimal set required for magnetosome biomineralization in MSR-1^36^, no magnetosomes have been detected in this organism. It might have several explanations: (i) the strain switches to the magnetotactic lifestyle only under very specific, yet not tested, conditions; (ii) it once was able to synthesize magnetosomes in its natural environment but had lost this ability upon subcultivation due to mutations before it was noticed; (iii) the strain does not naturally exploit magnetotaxis and its genes are non-functional or not actively expressed. To clarify which of these explanations is most likely, we first tested whether G2-11 can form magnetosomes under different laboratory conditions. To this end, the strain was cultivated photoheterotrophically, anaerobically or microoxically, in a complex medium with potassium lactate and soybean peptone, as commonly used for MSR-1 (FSM)^37^, as well as in minimal media with different C-sources previously shown to support growth in G2-11 (glucose, pyruvate, L-glutamine and ethanol)^26^. All media were supplied with 50 μM ferric citrate to provide sufficient iron for magnetite biomineralization. Since magnetosome biosynthesis is possible only under low oxygen tension, aerobic chemoheterotrophic growth of G2-11 was not tested. The best growth was observed in the complex FSM medium and a minimal medium with glucose or pyruvate, whereas L-glutamine and ethanol supported only weak growth (Supplementary Figure S2). Irrespective of the growth stage, none of the tested cultures demonstrated magnetic response as measured by a magnetically induced differential light scattering assay (Cmag)^38^. Consistently, micrographs of cells collected from the stationary phase cultures did not show any magnetosome-like particles (Supplementary Figure S2). This confirmed that G2-11 indeed cannot biosynthesise magnetosomes, at least under the conditions available for the laboratory tests. During cultivation, we also noticed that G2-11 cells did not move at any growth stage despite the initial description of this organism as motile using a single polar flagellum^26^, and containing several flagellum synthesis operons and other motility-related genes. Besides, the cells tended to adhere to glass surfaces under all tested conditions and form a dense clumpy biofilm immersed in a thick extracellular matrix (Supplementary Figure S2a-ii).

Considering that G2-11 generally lacks magnetosomes and appears to have a stationary lifestyle, which is not consistent with magnetotaxis, we assessed the maintenance of MGCs comes at fitness costs for the organism. To this end, we deleted the entire region containing the magnetosome genes (in the following, referred to as the MAI region) using the genetic tools we established for G2-11 in this work (Supplementary Figure S3a, see Materials and Methods for details). After the PCR screening, replica plating test, and genome re-sequencing, two clones of G2-11 ΔMAI were selected (Supplementary Figure S4). However, no significant differences were detected in the growth behaviour of the ΔMAI mutants compared to the wildtype (WT) control in a minimal medium supplied with acetate or pyruvate as a sole carbon source in two independent experiments (Supplementary Figure S3b). This finding suggests that the presence of the magnetosome genes neither provides benefits nor poses any substantial metabolic burden for G2-11, at least under the given experimental conditions.

### RNAseq reveals poor expression levels and antisense transcription in the MGCs of G2-11

Next, we set on to determine whether the magnetosome genes are transcribed in G2-11. For this, we analysed its whole transcriptome for the photoheterotrophic conditions, under which the best growth was observed, in two biological replicates. The expression levels of all the encoded genes calculated as TPM (transcripts per million) demonstrated a high correlation between the two replicates (Pearson’s correlation coefficient r = 0.98). Most genes of the *(mms6-like1)(mmsF-like1)mamH1IEKLMOH2* cluster were poorly or not transcribed at all (Fig. 2a, Supplementary data). Thus, transcription of *mms6-like1*, *mamF-like1*, *mamL, mamH1, mamI* and *mamK* did not pass the noise background threshold (TPM ≤ 2) in both replicates and were unlikely to be expressed, whereas *mamE, mamM, mamH2, feoAm,* and *feoBm* only slightly exceeded the threshold in at least one replicate and might be weakly transcribed (Fig. 2a). Although the TPM of *mamO* (TPM=5.67-6.10, Supplementary data) did exceed the threshold, the coverage plot reveals that the number of mapped reads sharply rises at its 3’-end, whereas the 5 ‘-end has low read coverage (Fig. 2b). This indicates the presence of an internal transcription start site (TSS) and its associated promoter within the coding sequence of *mamO* instead of the full transcription of the gene. Localization of an active promoter within *mamO* was recently described in MSR-1, suggesting that the transcriptional organization of MGCs may be more broadly conserved across MTB than assumed before^39^.

**Fig. 2.**
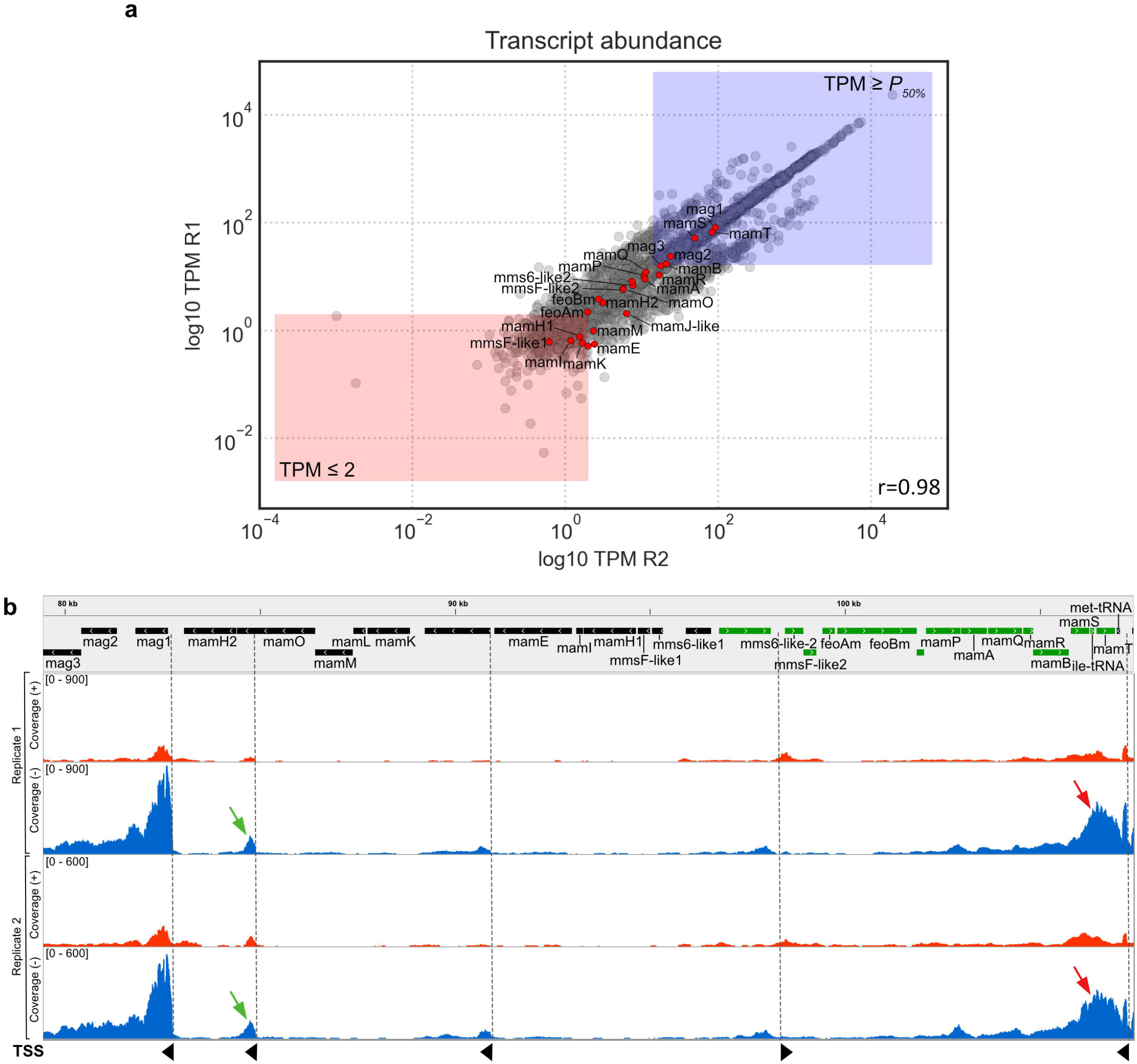
Transcription of the magnetosome genes in G2-11. (a) Log10 of the transcript abundances for all genes in the G2-11 genome presented as TPM (transcripts per million). Red dots represent the magnetosome genes. Red rectangle shows genes with TPM below the threshold, and blue rectangle shows genes with expression levels above median. R1 and R2: biological replicates. Pearson’s correlation coefficient (r) between the replicates is presented on the graph. (b) RNAseq coverage of reads mapped on the positive (red) and negative (blue) strands of the genome in the MAI region. The grey balk shows the gene map: genes encoded on the negative strand are coloured in black, on the positive - in green. Red arrows indicate the anti-sense transcription in the *mamPAQRBST* operon. Green arrows indicate the intragenic TSS within *mamO*. TSS are indicated with dashed lines and black arrowheads that show the direction of transcription.

Transcription of the genes within the *mag123*, (*mms6-like2*)(*mmsF-like2*), and *mamAPQRBST* clusters significantly exceeded the threshold, with the expression levels of *mag1, mamT* and *mamS* being above the overall median. At the same time, antisense transcription was detected in the *mamAPQRBST* region, with the coverage considerably exceeding the sense transcription (Fig. 2b).

This antisense RNA (asRNA) likely originated from a promoter controlling the tRNA gene positioned on the negative strand downstream of *mamT*. Such long asRNAs have the potential to interfere with the sense transcripts, thereby significantly decreasing expression of the genes encoded on the opposite strand^40^.

In summary, the RNAseq data revealed extremely low or lack of transcription of several genes that are known to be essential for magnetosome biosynthesis (*mamL, mamI, mamM, mamE* and *mamO*)^41,42^. Additionally, the detected antisense transcription can potentially attenuate expression of the *mamAPQRBST* cluster that also comprises essential genes, i.e., *mamQ* and *mamB*. Although other factors, like the absence of several accessory genes mentioned above and the potential accumulation of point mutations, might also be involved, the lack or insufficient transcription of the essential magnetosome genes appears to be the primary reason for the absence of magnetosome biosynthesis in G2-11.

### Magnetosome proteins from G2-11 are functional in a model magnetotactic bacterium

Although visual inspection of the magnetosome genes from the G2-11 did not reveal any frameshifts or other apparent mutations, accumulation of non-obvious functionally deleterious point substitutions in the essential genes could not be excluded. Therefore, we next tested whether at least some of the magnetosome genes from G2-11 still encode functional proteins that can complement isogenic mutants of the model magnetotactic bacterium MSR-1. In addition, we analysed the intracellular localization of their products in both MSR-1 and G2-11 by fluorescent labelling.

One of the key proteins for magnetosome biosynthesis in MSR-1 is MamB, as its deletion mutant is severely impaired in magnetosome vesicle formation and is entirely devoid of magnetite crystals^43,44^. Here, we observed that expression of MamB_[G2-11]_ partially restored magnetosome chain formation in MSR-1 Δ*mamB* (Fig. 3a, b-i, b-ii). Consistently, MamB_[G2-11]_tagged with mNeonGreen (MamB_[G2-11]_-mNG) was predominantly localized to magnetosome chains in MSR-1, suggesting that the magnetosome vesicle formation was likely restored to the WT levels (Fig. 3b-iii).

**Fig. 3.**
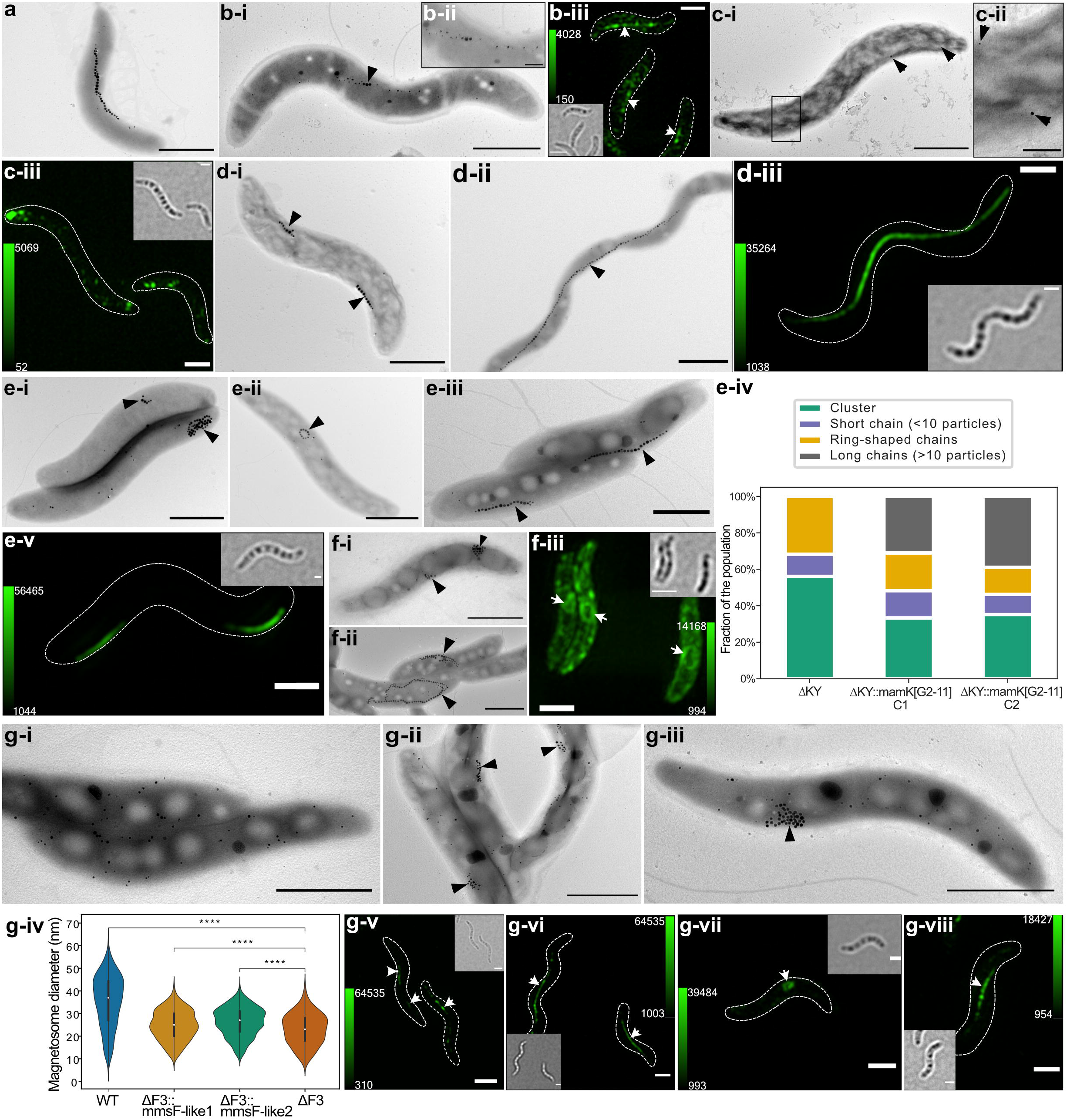
Genetic complementation and intracellular localization of magnetosome proteins from G2-11 in MSR-1 isogenic mutants. (a) TEM micrograph of MSR-1 wildtype (WT). (b) MSR-1 Δ*mamB*::*mamB*_[G2-11]_. (b-i) TEM micrograph and (b-ii) magnetosome chain close-up; (b-iii) 3D-SIM Z-stack maximum intensity projection of MSR-1 Δ*mamB*::*mamB*_[G2-11]_-mNG. (c) MSR-1Δ*mamQ::mamQ*_[G2-11]_. (c-i) TEM micrograph and (c-ii) close-up of the particles; (c-iii) 3D-SIM Z-stack maximum intensity projection. (d) MSR-1 Δ*mamK*::*mamK*_[G2-11]_. (d-i) TEM micrograph of MSR-1 Δ*mamK*; (d-ii) TEM micrograph of MSR-1 Δ*mamK*::*mamK*_[G2-11]_; (d-iii) 3D-SIM Z-stack maximum intensity projection of MSR-1 Δ*mamK*::mNG-*mamK*_[G2-11]_. (e) MSR-1 Δ*mamKY*::*mamK*_[G2-11]_. (e-i-ii) Representative cells of MSR-1 Δ*mamKY* mutant showing examples of a short chain, cluster (e-i), and ring-shaped chain (e-ii); (e-iii) TEM micrograph of MSR Δ*mamKY*::*mamK*_[G2-11]_ mutant showing the complemented phenotype; (e-iv) distribution of cells with different phenotypes in the populations of MSR-1 Δ*mamKY* and MSR-1 Δ*mamKY*::*mamK*_[G2-11]_ mutants (N > 50 cells for each strain population); (e-v) 3D-SIM Z-stack maximum intensity projection of MSR-1 Δ*mamKY*::mNG-*mamK*_[G2-11]_. (f) MSR-1 Δ*mamJ::mamJ*-like_[G2-11]_. (f-i) TEM micrograph of MSR-1 Δ*mamJ*; (f-ii) TEM micrograph of MSR-1 Λ*mam.J*::*mamJ*-like_[G2-11]_: (f-iii) 3D-SIM Z-stack maximum intensity projection of MSR-1 Δ*mamJ*::*mamJ*-like_[G2-11]_-*gfp*. (g) MSR-1 ΔF3::*mmsF-like1*_[G2-11]_ and ΔF3::*mmsF-like2*_[G2-11]_. (g-i) TEM micrograph of MSR-1 ΔF3; (g-ii) TEM micrograph of MSR-1 ΔF3::*mmsF*-like1_[G2-11]_; (g-iii) TEM micrograph of MSR-1 ΔF3::*mmsF*-like2_[G2-11]_; (g-iv) magnetosome diameter distribution in MSR-1 ΔF3 and the mutants complemented with *mmsF-* like1/*mmsF*-like2. Asterisks indicate points of significance calculated using Kruskal-Wallis test (**** indicates the p value < 0.0001); 3D-SIM Z-stack maximum intensity projections of: (g-v) MSR-1 ΔF3::*mNG-mmsF-like*1, (g-vi) MSR-1 WT::*mNG-mmsF-like*1, (g-vii) MSR-1 ΔF3::*mNG-mmsF-like2*_[G2-11]_, (g-viii) MSR-1 WT::*mNG-mmsF-like2*_[G2-11]_. Scale bars: all TEM micrographs, except close-up images, 1 μm; TEM close-ups, 0.2 μm; 3D-SIM, 1 μm. The calibration bars in 3D-SIM Z-stack projections indicate the minimum and maximum fluorescence intensity. Each 3D-SIM image is supplied with a bright field micrograph of the cells. Black and white arrowheads indicate magnetosomes in TEM and 3D SIM images, respectively.

Another essential protein MamQ is also involved in magnetosome vesicle formation, and its deletion eliminates magnetosomes in MSR-1 and other magnetospirilla^27,42^. Expression of MamQ_[G2-11]_ in MSR-1 Δ*mamQ* initiated the biosynthesis of very tiny and scarce magnetosomes (Fig. 3c-i, c-ii). mNG-MamQ_[G2-11]_ was localized in several intracellular patches, which distribution resembled that of the particles observed in the TEM micrographs (Fig. 3c-iii).

MamK is an actin-like filamentous protein, which is an essential structural component of the intracellular “magnetoskeleton” that aligns magnetosomes in linear chains^45^. Deletion of *mamK* in MSR-1 leads to the formation of disrupted short magnetosome chains instead of a continuous long chain typical for the WT (Fig. 3d-i). Expression of MamK_[G2-11]_ in MSR-1 Δ*mamK* resulted in the restoration of a normal magnetosome chain in most of the observed cells (Fig. 3d-ii). mNG-MamK_[G2-11]_ demonstrated linear signal indication the filament formation^46^ (Fig. 3d-iii). Since distinguishing Δ*mamK* from WT or a complemented phenotype can be difficult in shorter cells, we additionally transferred *mamK*_[G2-11]_ into the MSR-1 Δ*mamKY* mutant ^47^. In MSR-1 Δ*mamKY*, both magnetosome chains and their positioning are disrupted leading to the formation of magnetosome clusters or very short linear and ring-shaped chains (Fig. 3e-i, e-ii), which represent a more unambiguous phenotype than Δ*mamK*. Complementation of this mutant by a functional *mamK* should result in restoration of long chains, which would be positioned to the outer cellular curvature instead of the geodesic line of the helical cell since *mamY* is absent^47^. Indeed, expression of MamK_[G2-11]_ in MSR-1 Δ*mamKY* resulted in a population that included a considerable number of cells having long (≥10 particles) magnetosome chains, that were absent from MSR-1 Δ*mamKY* (Fig. 3e-iii). Evaluation of >50 cells for each of 2 randomly selected insertion mutants MSR-1 Δ*mamKY*::*mamK*_[G2-11]_ revealed that the long magnetosome chains were restored in 35-40% of the population (Fig. 3e-iv). Of note is that mNG-MamK_[G2-11]_ formed slightly shorter filaments in MSR-1 Δ*mamKY* than in Δ*mamK*, which were also characteristically displaced to the outer cell curvature due to the lack of *mamY*^47^ (Fig. 3e-v).

MamJ attaches magnetosomes to the MamK filament in MSR-1, mediating their chain-like arrangement. Elimination of *mamJ* disrupts this linkage, causing magnetosomes to aggregate owing to magnetic interactions^48^ (Fig. 3f-i). In MSR-1, MamJ is encoded within the *mamAB* operon, between *mamE* and *mamK*. Within the *(mms6-like1)(mmsF-like1) mamH1IEKLMOH2* cluster of G2-11, there is an open reading frame (ORF) encoding a hypothetical protein that is located in a syntenic locus (Fig. 1c). Although the hypothetical G2-11 protein and MSR MamJ differ considerably in length (563 vs. 426 aa), share only a low overall sequence similarity (31%), and are not identified as orthologues by reciprocal blast analyses, multiple sequence alignments revealed a few conserved amino acids at their N- and C-termini (Supplementary Figure S5). Moreover, in both proteins, these conserved residues are separated by a large region rich in acidic residues (pI 3.3 and 3.2) suggesting that the G2-11 protein might be a distant MamJ homolog. To test if it implements the same function as MamJ, we transferred this gene to MSR-1 Δ*mamJ*. Interestingly, it indeed restored chain-like magnetosome arrangement, which, however, often appeared as closed rings rather than linear chains (Fig. 3f-ii). Despite this difference, it indicated the ability of the hypothetical protein (hereafter referred to as MamJ-like _[G2-11]_) to attach magnetosomes to MamK, suggesting that in the native context, it can have a function identical to MamJ. Consistently, its fluorescently labelled version was often observed in ring-like structures within the cytoplasm of MSR-1 Δ*mamJ*, suggesting that it is indeed localized to magnetosomes (Fig. 3f-iii).

In magnetospirilla, magnetosome proteins MmsF, MamF and MmxF share an extensive similarity, and their individual and collective elimination gradually reduces the magnetite crystal size and disrupts the chain formation in MSR-1 (Fig. 3g-i; Uebe, manuscript in preparation). The MAI of G2-11 includes two genes, whose products have high similarity to these proteins, designated here as MmsF-like1_[G2-11]_ and MmsF-like2_[G2-11]_. Expression of each of them in the MSR-1 Δ*mmsF*Δ*mamF*Δ*mmxF* triple mutant (ΔF3) partially restored the magnetosome size that led to the formation of short magnetosome chains in MSR Δ*F3::mmsF-like1*_[G2-11]_ (Fig. 3g-ii) and clusters in MSR-1 ΔF3::*mmsF-like2*_[G2-11]_ (Fig. 3g-iii, iv). Consistently, fluorescently tagged mNG-MmsF-like1_[G2-11]_ and mNG-MmsF-like2_[G2-11]_ localized to magnetosomes in the pattern resembling that in the TEM micrographs of the complemented corresponding mutants (Fig. 3g-v, vii), or were perfectly targeted to the magnetosome chains in MSR-1 WT (Fig. 3g-vi, viii).

In G2-11, MamB_[G2-11]_-mNG, mNG-MamQ_[G2-11]_, MamJ-like_[G2-11]_-GFP, mNG-MmsF-like1_[G2-11]_ and mNG-MmsF-like2_[G2-11]_ were patchy-like or evenly distributed in the inner and intracellular membranes (Supplementary Figure S6). No linear structures that would indicate the formation of aligned magnetosome vesicles were observed in these mutants. As expected, mNG-MamK_[G2-11]_ formed filaments in G2-11 (Supplementary Figure S6c).

Expression of MamM, MamO, MamE, MamI and MamL failed to complement the corresponding deletion mutants of MSR-1 (not shown). Although detrimental mutations in the genes cannot be excluded, this result can be attributed to the lack of their native, cognate interaction partners, likely due to the large phylogenetic distances between the respective orthologues.

### Transfer of MGCs from MSR-1 endows G2-11 with magnetosome biosynthesis that is rapidly lost upon subcultivation

After having demonstrated the functionality of several G2-11 magnetosome genes in the MSR-1 background, we wondered whether, conversely, the G2-11 background is permissive for magnetosome biosynthesis. To this end, we transferred the well-studied MGCs from MSR-1 into G2-11, thereby mimicking an HGT event under the laboratory conditions. The magnetosome genes from MSR-1 were previously cloned on a single vector pTpsMAG1 to enable the one-step transfer and random insertion into the genomes of foreign organisms^23^. Three G2-11 mutants with different positions of the integrated magnetosome cassette were incubated under anaerobic phototrophic conditions with iron concentrations (50 μM) sufficient for biomineralization in the donor organism MSR-1. The obtained mutants indeed demonstrated detectable magnetic response (Cmag=0.38±0.11)^38^, and TEM confirmed the presence of numerous electron-dense particles within the cells (Fig. 4), which, however, were significantly smaller than the magnetosome crystals of MSR-1 (18.5±4.3 nm to 19.9±5.0 nm vs 35.4±11.5 nm in MSR-1, Fig. 4b) and formed only short chains or were scattered throughout the cells (Fig. 4a, c-i). Mapping of the particle elemental compositions with energy-dispersive X-ray spectroscopy (EDS) in STEM mode revealed iron- and oxygen-dominated compositions, suggesting they were iron oxides. High-resolution TEM (HRTEM) images and their FFT (Fast Fourier Transform) patterns were consistent with the structure of magnetite (Fig. 4c). Thus, G2-11 was capable of genuine magnetosome formation and even occasionally built magnetosome chains after acquisition of the MGCs from MSR-1.

**Fig. 4.**
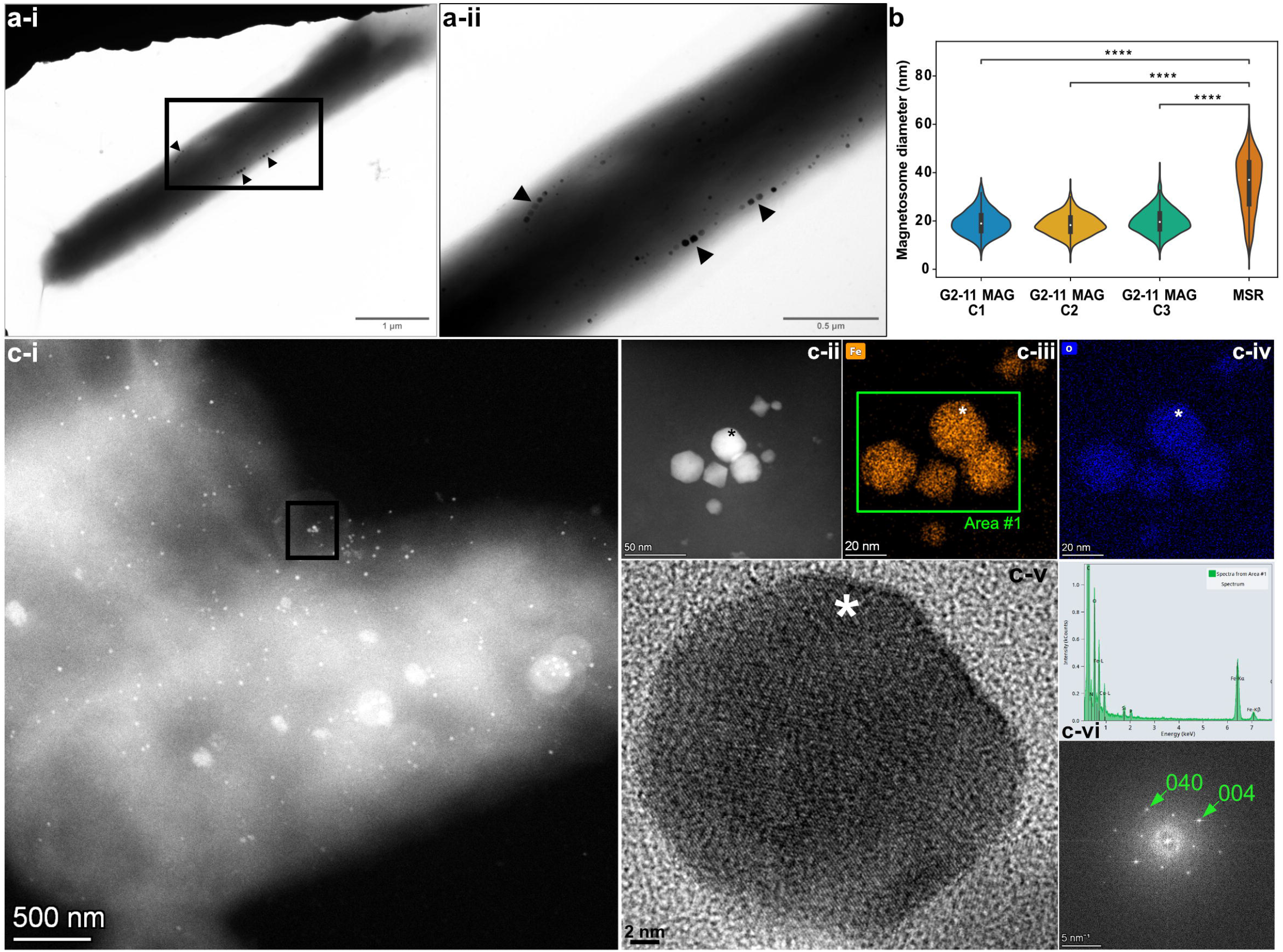
Magnetosome biosynthesis by G2-11 upon transfer of the MGCs from MSR-1. (a) A cell with magnetosomes (i) and a close-up of the area with magnetosome chains (ii). (b) Violin plots displaying magnetosome diameter in three MAG insertion mutants of G2-11 in comparison to MSR-1. Asterisks indicate points of significance calculated using the Kruskal-Wallis test (**** designates p-value < 0.0001). (c) Crystallography analysis of magnetosomes from G2-11 MAG: (c-i) HAADF image of a cell; (c-ii) HAADF image of the cluster from the area shown with a black frame in (c-i); (c-iii) iron (Fe) and (c-iv) oxygen (O) EDS elemental maps of the magnetosome cluster. The peak indicating Cu is an artefact from the copper grid; (c-v) HRTEM image of the magnetosome crystal marked with an asterisk in (cii-iv); (c-vi) EDS spectrum from Area #1 in c-iii; (c-vii) FFT pattern corresponding to the HRTEM in c-v, obtained along the [100] axis of magnetite.

In at least three independent transfer experiments, we noticed that the ability to synthesize heterologous magnetosomes was highly unstable in G2-11 upon subcultivation. The Cmag of the transgenic cultures started to decline soon after the transfer, and the magnetic response became eventually undetectable in all of them after 10-15 daily culture passages. Concurrently, the mutant cells in this non-magnetic state were devoid of magnetosomes. To understand the mechanism of the trait loss, we sequenced the genomes of three randomly selected newly “magnetized” mutants (hereafter, C1-3) immediately after the genetic transfer, and again after the magnetic response had been lost from the cultures. All three mutants demonstrated a rapid decline in Cmag after the 8^th^ passage (Fig. 5a), whereas the rate at which the cultures transitioned to a completely non-magnetic state (nonMAG) varied among the clones. As expected, TEM observations confirmed the loss of magnetosomes (Fig. 5b). Genome analysis showed that in two out of three insertion mutants (C2 and C3), the entire integrated magnetosome cassette was deleted in their nonMAG descendants (Fig. 5c). Visual inspection of the reads mapped to the insertion locus and the sequences flanking the integrated cassette revealed that a large fraction of the reads (87.4% and 96.9% in C2 and C3, respectively) was mapped to a restored wildtype sequence (except leaving a single nucleotide insertion in place of the deleted cassette), indicating a complete excision of the integrated cassette in most cells (Fig. 5d). Since on pTpsMAG1 the magnetosome cassette is flanked by inverted repeats recognised by the mariner transposase for mobilisation and insertion, we believe that these repeats could be recognised and re-used for the excision of the cassette in G2-11, mediated either by intrinsic recombinases or one of many transposases encoded in its genome.

**Fig. 5.**
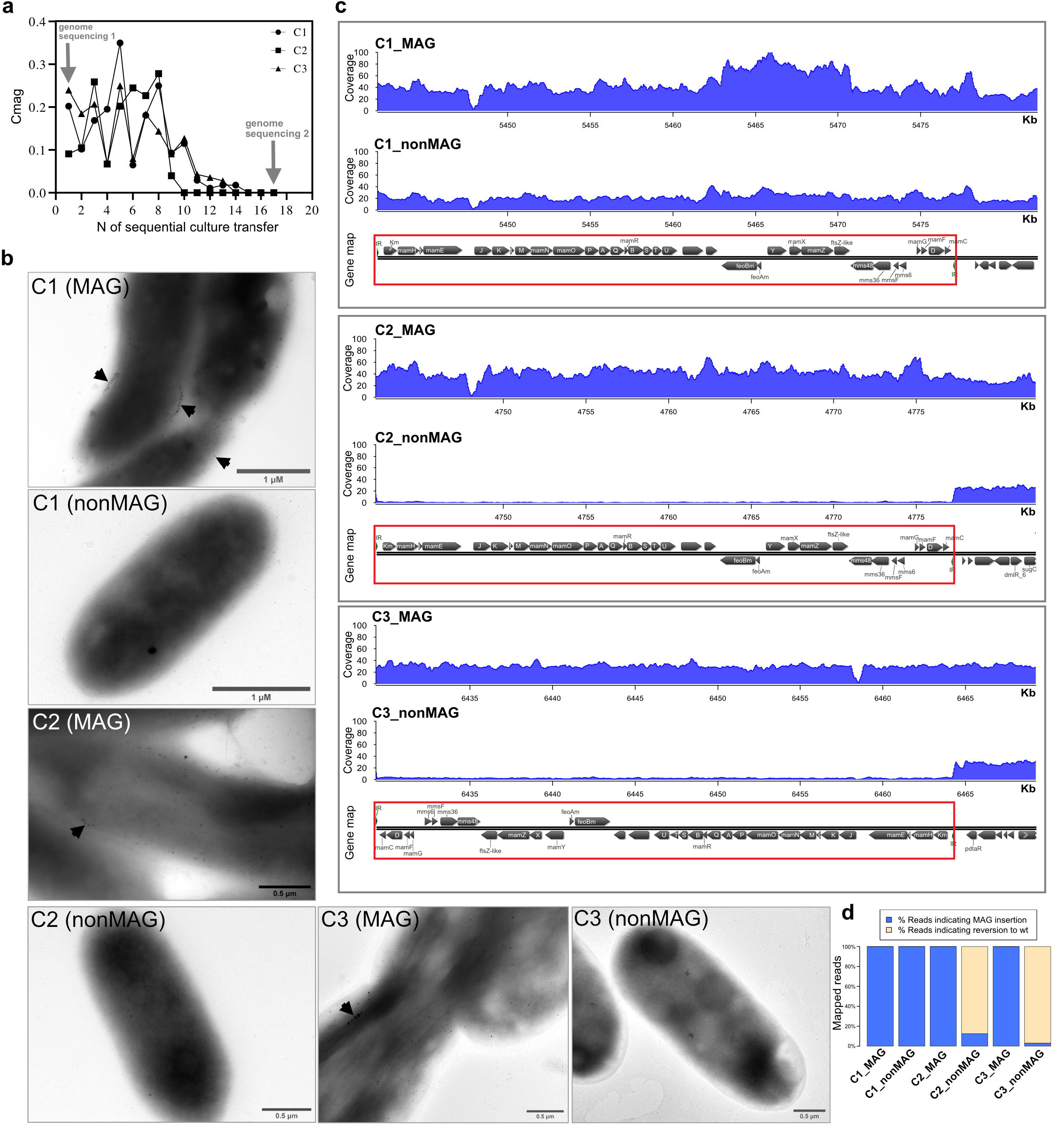
Analysis of the dynamics of magnetosome biosynthesis loss in G2-11 MAG. (a) Change of the magnetic response (Cmag) of three randomly selected MAG insertion mutants with sequential culture passages. Arrows indicate the timepoints at which the genomes were re-sequenced. (b) TEM micrographs of the cells at timepoint 1 (MAG) and timepoint 17, after the loss of magnetosomes (nonMAG). (c) Read coverage normalized to the library size of the MAG and nonMAG mutants. The gene maps show the insertion positions of the MAG cassette (highlighted by red rectangle) within the genome. (d) Bar chart showing the percentage of reads indicating the MAG cassette excision and the cassette presence.

In contrast to C2 and C3, no mutations could be detected in the nonMAG state of C1. This suggests that, in addition to the cassette deletion, other mechanisms to suppress the expression of foreign magnetosome genes, e.g., transcriptional silencing, are likely involved. Besides, the native MGCs present in G2-11 were not affected in either of the mutants. Overall, this experiment demonstrated that although G2-11 can synthesize magnetosomes upon acquisition of the foreign magnetosome genes, their expression imposes a significant negative selection pressure on G2-11, causing the gene deletion (C2 and C3) or potential suppression of expression (in C1).

## Discussion

In the current study, we present the discovery and comprehensive analysis of MGCs in the non-magnetotactic phototrophic bacterium *R. atsumiense* G2-11. Compared to many well-known silent biosynthetic clusters that control secondary metabolite production pathways^49,50^, this is, to our knowledge, the first evidence for the existence of silent genes encoding the biosynthesis of magnetic organelles in non-MTB. Our analyses suggest a foreign origin of the MGCs in G2-11, deriving them from an alphaproteobacterial MTB likely belonging to *Azospirillaceae*. The compact organization, tetranucleotide usage bias and the fact that no MTBs have been revealed among *Acetobacteraceae* to date support a rather recent HGT of the magnetosome genes to G2-11. Despite the lack of other known representatives of the genus *Rhodovastum*, it can be estimated that the HGT event took place at least after the delineation of the contemporary extant *Acetobacteraceae* genera. These data allow us to speculate that HGT of magnetosome genes occurs more frequently than suggested before^9^ and that organisms like G2-11 can contribute to spreading and evolution of MGCs.

Why were the magnetosome genes not eliminated in G2-11 despite not conferring any selective advantage to the host^51^? Several of our findings provide a likely explanation. First, approximately half of the genes within the MGCs of G2-11 are not or only weakly transcribed, whereas expression of the second half is likely attenuated by asRNA^40^. Hence, we hypothesise that their maintenance within the genome might be neutral to the strain fitness. Indeed, a comparison of the G2-11 ΔMAI mutant and the WT revealed no significant effect of the magnetosome gene presence on the strain growth, at least under the experimental conditions tested, which essentially places the MGCs of G2-11 under neutral selection.

By complementation of isogenic mutants, we demonstrate that magnetosome genes *mamB, mamQ, mamK, mamJ-like, mmsF-like1*, and *mmsF-like2* from G2-11 are functional in MSR-1 and their products localize to magnetosomes. The fact that other genes previously shown to be essential in MSR-1, i.e., *mamM, mamO, mamE, mamI* and *mamL*, failed to restore the magnetic phenotype in the MSR-1 mutants might be attributed to the protein malfunction in a foreign context, as magnetosome proteins are known to rely on a tight interaction network with other co-evolved magnetosome proteins^43,52^. Furthermore, the lack of complementation in MSR-1 for some magnetosome genes from distantly related MTB was observed previously^43^.

Finally, we tackled the question of whether G2-11 is generally capable of magnetosome biosynthesis if provided with a complete and functional gene set from a foreign donor. Intriguingly, the obtained G2-11 mutants were able to produce numerous small magnetosomes, occasionally assembled into short chains. This indicates that G2-11 provides an appropriate and ‘permissive’ auxiliary genetic background that supports magnetosome formation upon acquisition of the MGCs from an MTB. However, unlike previously synthetically ‘magnetised’ foreign hosts, in which the phenotype was stable^22,23^, G2-11 reproducibly experienced the rapid loss of the magnetic phenotype due to extensive deletions or potential silencing. This finding implies that functional expression of the acquired MGC poses a considerable metabolic burden on G2-11, which stimulates the use of various mechanisms to either avoid unwanted expression or eliminate the genes. These results shed light on the potential state of the G2-11 cells immediately after the HGT of its ‘native’ magnetosome genes: in the absence of conditions selecting for the magnetosome presence, the phenotype was quickly lost, likely due to the inconsistency between the conferred function and the host’s stationary lifestyle.

In summary, this is the first reported case of a natural occurrence of MGC in a photosynthetic non-magnetotactic bacterium, which reveals that transcriptional inactivation may serve as an important mechanism for preserving genes encoding a complex trait such as prokaryotic organelle biosynthesis in a latent state within genomes. Although these MGCs are currently dormant, they may serve as a “raw material” for quick adaptation to changing environmental conditions or new niche colonisation, giving this organism the potential to evolve into an MTB. Besides, G2-11 provides a rare glimpse into the mostly hidden world of genetic exchange of large gene clusters between microbes. Thus, the organisms that preserve such clusters in a silent but functional state may arguably serve as “transshipment points” for further gene transfer, with a potential to promote their expansion in native communities.

## Materials and Methods

### Strains and conditions

*Rhodovastum atsumiense* strain G2-11 (DSM 21279) and *Magnetospirillum gryphiswaldense* strain MSR-1 (DSM 6361) were routinely cultivated in flask standard medium (FSM, 10 mM HEPES [pH 7.0], 15 mM potassium lactate, 4 mM NaNO3, 0.74 mM KH2PO4, 0.6 mM MgSO47H2O, 50 mM ferric citrate, 3 g/liter soy peptone, 0.1 g/liter yeast extract), in 15- or 50-mL screw-capped falcon tubes filled to 3/4 of their volume, at 120 rpm agitation. For phototrophic growth, G2-11 was cultivated in Hungate tubes filled with medium to 2/3 of their volume with the headspace containing 100% N_2_, at a light intensity of 50 μmol/s/m^2^, without agitation. For the test of different conditions for magnetosome formation and the growth comparisons, G2-11 was cultivated in the minimal medium consisting of the mineral base RH2 supplemented with 15 mM of the organic carbon source, 50 μM ferric citrate, and 0.01% yeast extract. Selection for the mutants was carried out on FSM solidified with 1.5% (wt/vol) agar and supplemented with antibiotics: 5 μg/mL of kanamycin (Km) for MSR-1 and G2-11, 6 μg/mL of tetracycline (Tc) for G2-11, 20 μg/mL of Tc for MSR-1, 5 μg/mL of gentamycin (Gm) for G2-11.

*E. coli* WM3064 strains carrying plasmids were cultivated in lysogeny broth (LB) supplemented with 0.1 mM DL-α,ε-diaminopimelic acid (DAP) and 25 μg/ml Km, 12 μg/ml Tc or 15 μg/ml of Gm at 37°C, with 180 rpm agitation. Characteristics of the strains used in the study are summarized in Supplementary Table S2.

### Genome sequencing and assembly

A closed reference genome of *Rhodovastum atsumiense* strain G2-11T (DSM 21279) was obtained from the genomic DNA extracted from 2-10 mL of stationary cultures with the Zymo Research Midiprep gDNA kit. Sequencing and assembly was performed mixing Illumina technology and Oxford Nanopore technology (ONT). First, for Illumina sequencing, 250 ng DNA was sonicated to a 100–1,000 bp size range using the E220 Covaris Focused-Ultrasonicator (Covaris, Inc.). The fragments were end-repaired, then 3’-adenylated and NEXTflex HT Barcodes (Bio Scientific Corporation) were added using NEBNext DNA modules products (New England Biolabs). After two consecutive cleanups with 1×AMPure XP, the ligated product was amplified by 12 PCR cycles using the Kapa Hifi Hotstart NGS library amplification kit (Kapa Biosystems), followed by purification with 0.6×AMPure XP. After library profile analysis conducted by an Agilent 2100 Bioanalyzer (Agilent Technologies) and qPCR quantification (MxPro, Agilent Technologies), the library was sequenced on an Illumina MiSeq with a MiSeq Reagent Kit v.2 (2 × 250 bp; Illumina Inc.). A total of 3.27 x 106 paired-end reads were obtained. The Illumina reads were trimmed by removing low-quality nucleotides (Q < 20), sequencing adaptors and primer sequences using an internal software based on the FastX package (http://hannonlab.cshl.edu/fastx_toolkit/index.html). Reads shorter than 30 nucleotides after trimming were also discarded. For Nanopore sequencing, library preparation was done with 1 μg of the same input DNA following the 1D Native barcoding genomic DNA protocol with EXP-NBD104 and the SQK-LSK109 ligation kit (Oxford Nanopore). The library was sequenced using a Nanopore R9.4.1.revD flow cell (Oxford Nanopore) and the PromethION device with the MinKNOW v.4.0.5 and Guppy v.4.0.11 software. A total of 309 928 reads were obtained with a N50 of 8.07 Kb. Two hybrid assemblies were launched in parallel with Unicycler v.0.4.6 (default options) and Unicycler v.0.4.6 (—sc option for SPAdes) and compared to circularize chromosomes and plasmids^53^. The final assembly resulted in a single chromosome 6.48 Mb in length with a GC content of 68.96% and 8 plasmids (from 220 Kb to 10.69 Kb) with an average GC content of 66.07%. Assembly completeness and contamination were estimated at 99.5 and 0.54%, respectively, using checkM v.1.0.11^54^ with a set of 336 *Rhodospirillales-specific* markers. The automatic annotation was performed with the MicroScope platform (https://mage.genoscope.cns.fr/microscope)^55^.

To test the ‘magnetized’ mutants for the presence or absence of the MAG cassette, genome sequencing was performed again. To this end the gDNA was extracted as described above. Library preparation and sequencing were performed at Novogene (UK) Co. Ltd using Illumina NovaSeq 6000 with paired-end 2 × □ 150 □ bp reads corresponding to 1.0-1.3 Gbp in different samples (estimated coverage 133×). For the raw reads produced with Illumina, adapter trimming and quality control filtering were carried out with *fastp* using standard parameters^56^. Processed reads were mapped to the reference genome using *Bowtie 2*^57^ and Geneious (8.1.9)^58^. The read coverage for gDNA re-sequencing in magnetized mutants was calculated using the *bamCoverage* tool, a part of *deepTools* (v. 3.3.2) programs set^59^ available on the Galaxy server (https://usegalaxy.eu)^60^. For this, the number of reads that overlap 50 nt bin fragments in the genome was counted and normalized to the number of mapped reads per million. The resulting binned counts per million (CPM) were processed as *bedgraph* or *bigwig* format files. The coverage histograms were visualized using R package *Sushi* v.1.32.0 (Phanstiel, D. H. Sushi: Tools for visualizing genomics data. R package version 1.16.0. https://www.bioconductor.org/packages/release/bioc/html/Sushi.html (2019)). The fraction of reads indicating reversion to the wildtype locus by excision of the MAG cassette was calculated after mapping reads to the wildtype and the mutant reference genomes by variant calling implemented in Geneious (8.1.9) and checked by visual inspection

### RNA sequencing

RNA sequencing was performed for two biological replicates. For RNA isolation, 200 mL of each replicate culture was grown in 250-mL screw-cap bottles at 28 °C, 180 rpm agitation, with incandescent light (~1500 lux). Cells were harvested at mid-logarithmic phase (optical density at 660 nm [OD660] = 0.130 and 0.159 for replicate 1 and 2, respectively) by centrifugation at 4000 × rpm for 10 min at 4 °C using an Allegra^®^ X-15R centrifuge (Beckman Coulter) and flash-frozen in liquid nitrogen before RNA isolation. Total RNA was extracted using hot phenol method^61^ with modifications. Briefly, cell pellets were resuspended in 2.5 mL of an ice-cold solution containing 0.3M sucrose and 0.01M sodium acetate (pH 4.5). Lysis occurred by careful mixing the resuspended cells with 2.5 mL of hot (65 °C) solution containing 2% SDS and 0.01M sodium acetate (pH 4.5). The equal volume of hot (65 °C) phenol was added to the lysed cells and mixed by inverting. Tubes were briefly chilled in liquid nitrogen and centrifuged at 4,700 rpm for 5 min, 4 °C. The aqueous layer was used for sequential re-extraction with 5 mL of hot (65 °C) phenol, 5 mL of phenol:chloroform:isoamyl alcohol (in proportion 25:24:1, pH 4.5), and chloroform:isoamyl alcohol (24:1). RNA was precipitated by incubation with 2.5 volume of 100% ethanol and 0.1 volume of 3M sodium acetate (pH 4.5) at −80 °C for 30 min. RNA was pelleted by centrifugation at 4,700 rpm for 30 min, 4 °C, washed twice with ice-cold 75% ethanol. After drying in the air, the pellet was resuspended in 200 μL of RNAase-free water. The RNA concentration and quality were controlled by Nanodrop measurements, electrophoresis in 1% agarose gels and the Agilent 2100 Bioanalyzer. Library preparation and RNA sequencing was carried out by Novogene Ltd. (UK). Before library construction, mRNA was enriched using oligo(dT) beads and rRNA was removed using the Ribo-Zero™ kit. The strain-specific cDNA library was prepared with an Illumina kit according to the manufacturer’s recommendation and sequenced using the Illumina Novaseq 6000. The RNA-seq reads were mapped to the genome using HISAT^64^ aligning program with strand-specific parameters. The resulting alignments and the reference annotation were used for *de novo* transcript assembly and prediction of transcription levels with StringTie^65^. The transcription levels were calculated as transcripts per million (TPM)^66^. A threshold of TPM≥2 was applied to define the expressed genes^67^.

### Molecular phylogeny

Since the previously built tree based on the 16S rRNA gene affiliated *Rhodovastum atsumiense* to the phylum *Alphaproteobacteria*, order *Rhodospirillales*, family *Acetobacteraceae*^26^, we investigated the evolutionary relationships between strain G2-11 and its *Acetobacteraceae* relatives using whole genome sequences. First the proteome of all non-redundant (1 representative species per genus) closed genomes of *Acetobacteraceae* (LPSN nomenclature, ranked at the order level in GTDB) were downloaded from the public repository RefSeq database at NCBI in April 2022. This database was enriched with the draft genome of the closest relatives of *R. atsumiense: Acidisphaera rubrifaciens* strain HS-AP3 and *Rhodopila globiformis* strain DSM 161. The dataset was also completed with a set of genomes from the *Rhodospirillaceae* family and the *Azospirilllaceae* family that was then used as an external outgroup based on the latest *Alphaproteobacteria* phylogeny^29^. A set of 54 proteins composing the 30S and 50S ribosome subunits (RPs) were extracted using HMM profiles^68^, while the magnetosome proteins sequences were retrieved using the Microscope annotation pipeline. Shared magnetosome proteins MamKMOPAQBST were further used to infer the origin of G2-11 MGC. For this, complete genomes of the closest non-alphaproteobacterial MTB were used as an external group.

For both ribosomal proteins and magnetosome proteins-based trees, amino acids sequences were aligned independently using MAFFT version 7.490^69^ and alignments were trimmed using BMGE^70^ setting block length and gap frequency to 3 and 0.5 respectively, and using the BLOSUM30 matrix. Maximum-Likelihood tree was built from the concatenated sequences with IQ-TREE version 2.1.4^71^ and a partition model. For that purpose, a model selection was performed on each protein sequence alignment in the concatenation. Branch support was estimated through non-parametric bootstrap with 500 replicates.

### Molecular and genetic methods

Oligonucleotides applied in this study are listed in Supplementary Table S3. For the complementation experiments, magnetosome genes were PCR amplified from the G2-11 genome using the high-fidelity Q5® polymerase (New England Biolabs, New England USA) and cloned by restriction sites into the pBamII-Tc vector (Supplementary Table S4). To analyse the intracellular localization, the proteins that showed complementation were N- or C-terminally tagged with mNeonGreen (mNG)^72^ or GFP that were codon-optimized for MSR-1 and expressed under the control of P_mamDC45_ promoter^73^. For this, the genes were fused to the fluorescent protein gene separated by a GSA linker and cloned into the pBamII-Tc vector using Gibson assembly^74^. For the transfer of magnetosome gene clusters from MSR-1 to G2-11 the previously constructed vector pTpsMAG1 was applied^23^.

The plasmids were transferred into MSR-1 and G2-11 by biparental conjugation as described elsewhere^75^, with the following modifications for G2-11: the conjugation mixture was inoculated on several selection plates with 10^-2^ to 10^-4^ dilutions, which were incubated for 5-7 days aerobically in the dark, at 28 °C.

### Generation of the ΔMAI mutant of G2-11

To assemble the vectors for deletion of the MAI region in G2-11, the 1.2 kb left (LHR) and right flanking homology regions (RHR) were PCR amplified using specific primers. The LHR was cloned into the vector pAL01-MCS1 -Km^36^ by *KpnI* and *NotI* restriction sites, the RHR was cloned into pAL02-MCS2-Gm^36^ by *KpnI* and *BamHI* (Supplementary Figure S3). Two versions of Cre-expressing vector pCM157 were tested separately for the region excision: (i) with *cre* gene under the control of inducible *P_lac_* promoter^76^, and (ii) under the control of *P_nir_* promoter from MSR-1^77^. In the culture bearing pCM157-P*lac* expression of Cre was induced by adding 2 mM and 4 mM IPTG, and with pCM157-P_*nir*_ by incubation of the culture under anaerobic phototrophic conditions. The deletion protocol was applied as described previously^76^ with the following modifications: the cultures were incubated for 6 days under the Cre-expressing conditions with a daily check for the desired mutation by PCR screening. After the band appeared, the culture was plated with dilutions 10^-4^ to 10^-7^. The colonies were scaled-up without antibiotics and checked using replica plating with and without antibiotics for the excision of the integrated plasmids and curing of pCM157. The desired mutation was confirmed by PCR and subsequent genome sequencing using the Illumina NovaSeq 6000 as described previously.

### Fluorescence 3D-SIM microscopy

For fluorescence microscopy, 2 μL of the cells were immobilized on 1% (wt/vol) agarose pads. 3D-SIM (striped illumination at 3 angles and 5 phases) was performed on an Eclipse Ti2-E N-SIM E fluorescence microscope (Nikon) equipped with a CFI SR Apo TIRF AC 100×H NA1.49 Oil objective lens, a hardware-based ‘perfect focus system’ (Nikon), LU-N3-SIM laser unit (488/561/640 nm wavelength lasers) (Nikon), and an Orca Flash4.0 LT Plus 17 sCMOS camera (Hamamatsu). 3D-SIM z-series for *R. atsumiense* were acquired at a total thickness of −1.44 to 1.44 μm with 0.12 μm z-step spacing and for MSR-1 at a total thickness of −1.25 to 0.75 μm with 0.12 μm z-step spacing. The exposure time was in the range of 20 to 300 ms at 60% laser power. Fluorescence excitation of mNG was at 403.5 nm and emission was detected at 522.5 nm. Image reconstruction was performed in NIS-Elements 5.01 (Nikon) using the ‘stack reconstruction’ algorithm with the following parameters. The ‘illumination modulation contrast’ was set to ‘auto’. The ‘high-resolution noise suppression’ was set to 0.1^78^. Images were analysed using ImageJ 1.53c^79^.

### Transmission electron microscopy (TEM) and HRTEM

Samples for conventional TEM were concentrated from 2-3 mL cultures by centrifugation, adsorbed on carbon-coated copper grids and imaged using a JEOL-1400 Plus or JEOL-2100 TEM (Japan) at 80 kV acceleration.

High-resolution (HRTEM) images and selected area electron diffraction (SAED) patterns were obtained using the TEM mode of a Talos F200X G2 instrument (Thermo Fisher Scientific, Waltham, MA, USA) at 200 kV accelerating voltage. The same device was used for scanning transmission microscopy (STEM) high-angle annular dark-field (HAADF) images that were collected for both imaging and mapping elemental compositions by coupling the STEM mode with energy-dispersive X-ray spectrometry (EDS).

### Statistical analysis and data visualization

Chromosome and plasmid structures were visualized using CGView 1.7 package^80^. Statistical analysis and graph plotting were carried out using in-house scripts written in Python v. 3.8. Analysis and visualization of the transcription data were conducted using python libraries *pandas 1.0.3*^81^ and *matplotlib 3.5.1*^82^. Tetranucleotide-derived z-scores of the MAI and the reference regions were calculated as described in Teeling et al. (2004)^62^ using *pyani* (v. 0.2.11) package^63^ and a custom script. The magnetosome diameters were measured using imageJ v.1.53c, and the violin plots were constructed applying libraries *seaborn 0.112*^83^ and *matplotlib*. The Kruskal-Wallis analysis was conducted using the library *statannot* (https://github.com/webermarcolivier/statannot).

## Data availability statement

Sequencing reads and the annotated complete genome of strain *Rhodovastum atsumiense* strain G2-11 (DSM 21279) wildtype was deposited to the European Nucleotide Archive database under the BioProject number PRJEB52102. RNA-seq reads and sequencing reads for the genomes of G2-11 mutants generated in this study were deposited to NCBI GenBank database under the BioProject number PRJNA818516.

## Acknowledgments

This study was supported by the European Research Council (ERC) under the European Union’s Horizon 2020 research and innovation program (Grant No. 692637 to D.S.), the Deutsche Forschungsgesellschaft (DFG) (Grant UE 200/1-1 to RU), and ANCESMAG ANR project (ANR-20-CE92-0050).

We thank Corinne Cruaud and the Genoscope (CEA, Commissariat à l’Energie Atomique et aux Energies Alternatives) at Evry (France) for their essential contribution to the efficiency of the platform.

The Galaxy server that was used for some calculations is in part funded by Collaborative Research Centre 992 Medical Epigenetics (DFG grant SFB 992/1 2012) and German Federal Ministry of Education and Research (BMBF grants 031 A538A/A538C RBC, 031L0101B/031L0101C de.NBI-epi, 031L0106 de.STAIR (de.NBI)). Electron microscopy at Nanolab, University of Pannonia was supported by the National Research, Development and Innovation Office, Hungary (grant NKFIH-471-3/2021). We are grateful to M. Schüler and S. Geimer for their help with electron microscopy. We also thank J. Kachel and U. Brandauer for technical assistance.

## Author contributions

R. U. and M.V.D. conceived the study and designed the experiments. M.V.D., A.P., L.S., and R.P.A. conducted the experiments; M.P. carried out the HRTEM and crystallography analysis; C.L.M. and S. F. sequenced and assembled the complete genome; C.L.M. contributed to phylogenetic analysis. M.D., R.U. and D.S. wrote the manuscript. All authors read and approved the final manuscript.

## List of supplementary materials

Supplementary Figure S1

Supplementary Figure S2

Supplementary Figure S3

Supplementary Figure S4

Supplementary Figure S5

Supplementary Figure S6

Supplementary data(.csv)

Supplementary Table S1

Supplementary Table S2

Supplementary Table S3

Supplementary Table S4

## Supplementary figure captions

Supplementary Figure S1 Plasmids of *Rhodovastum atsumiense* G2-11

Supplementary Figure S2 Photoheterotrophic cultures (i) and TEM micrographs (ii) of G2-11 cells grown under phototrophic conditions with various carbon sources: (a) complex medium with potassium lactate and peptone (FSM); (b) minimal medium with glucose; (c) minimal medium with pyruvate; (d) minimal medium with L-glutamine; (e) minimal medium with ethanol. Black arrowheads: bacteria adherent to the glass surface; red arrowhead: a matrix encompassing cell clumps; PP: putative polyphosphates; PHB: putative polyhydroxybutyrate inclusions.

Supplementary Figure S3 Construction scheme and growth analysis of the G2-11 ΔMAI mutants. (a) The MAI region was excised using a Cre-lox-based allele exchange technique with modifications (see “Materials and Methods” for details). LHR: left homologous region; RHR: right homologous region. (b) Growth curves of *two* G2-11 ΔMAI clones and the wildtype (WT) cultivated chemoheterotrophically in a minimal medium with acetate or pyruvate as carbon source. Results of two independent experiments are shown (i and ii). Error bars indicate the standard deviation of three biological replicates.

Supplementary Figure S4 Screening for G2-11 ΔMAI mutants. (a) PCR-screening of the transconjugants: (i) screening of the culture for the band indicating deletion in the clones with pCM157-P_lac_ (left) and pCM157-P_nir_ (right). (ii) Screening of the individual clones for (left to right): the band indicating deletion, the excision of the pAL plasmids from the flanking regions. (iii) Test for the presence of the magnetosome genes in the genome of the candidate ΔMAI clones. The clones selected for further work are highlighted by yellow ellipses. (b) Replica-plating of the clones on agar plates with and without antibiotics. The selected mutants are highlighted in red punctuated circles.

Supplementary Figure S5 Alignment of MamJ-like protein sequences from MTB and G2-11. The fragments that show sequence conservation are shown in close-ups. Note that the gene that occupies the syntenic locus of *mamJ* in CCP-1 is dissimilar to both MamJ-like from G2-11 and MamJ proteins of magnetospirilla.

Supplementary Figure S6 3D-SIM Z-stack maximum intensity projection of G2-11 WT expressing magnetosomes proteins tagged by mNeonGreen: (a) WT::*mamB*_[G2-11]_-*mNG*; (b) WT::*mNG-mamQ*_[G2-11]_; (c) WT::*mNG-mamK*_[G2-11]_, (d) WT::*mamJ-like*_[G2-11]_-gfp; (e) WT::*mNG-mmsF-like1*_[G2-11]_, (g) WT::*mNG-mmsF-like2*_[G2-11]_. Scale bars: (a-d), 1 μm; (e),(g), 2 μm.

