## Supplementary figures and images for "Hidden Talents: Silent Gene Clusters Encoding Magnetic Organelle Biosynthesis in a Non-Magnetotactic Phototrophic Bacterium"

### Supplementary Figure S1

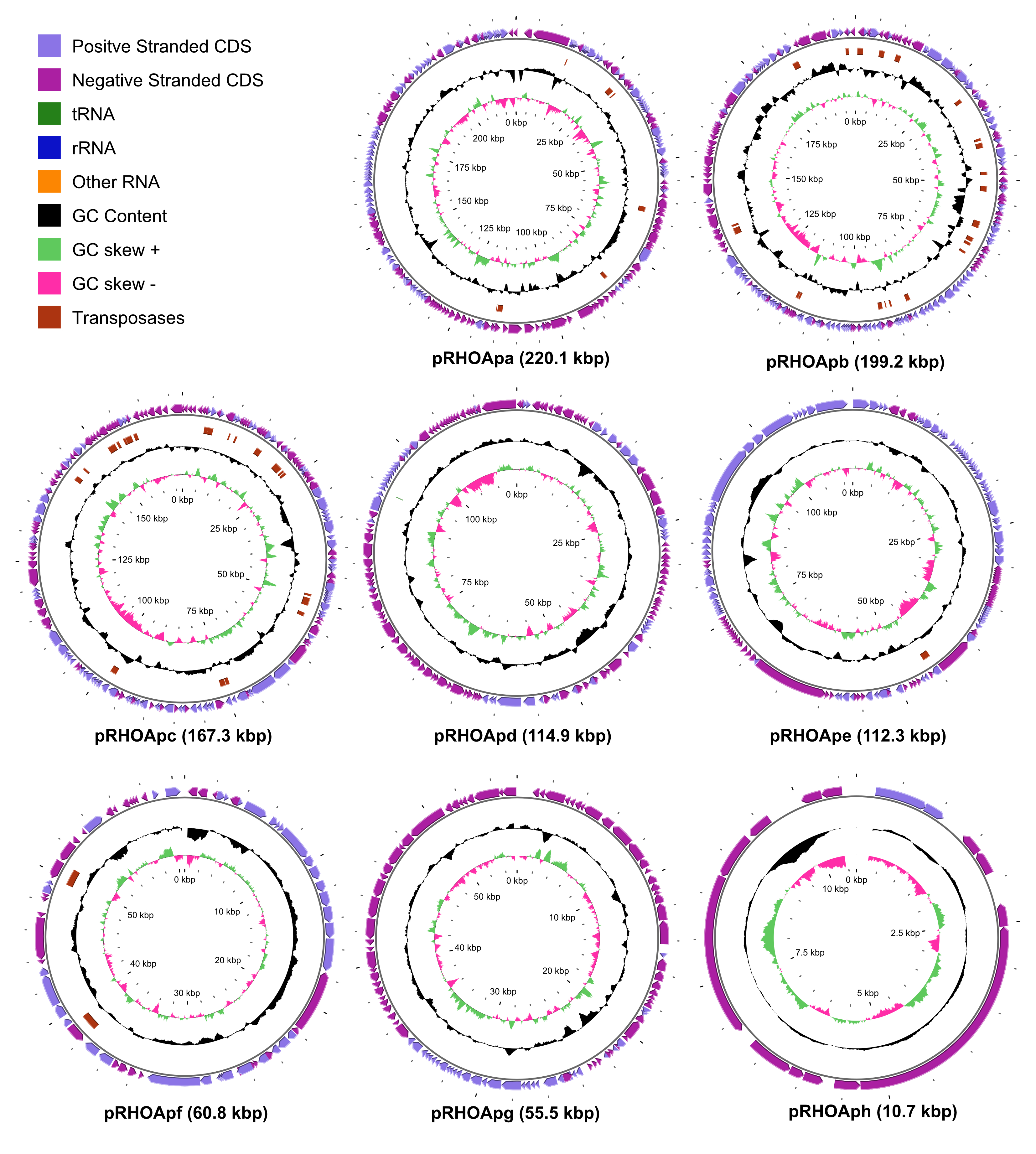

### Supplementary Figure S2

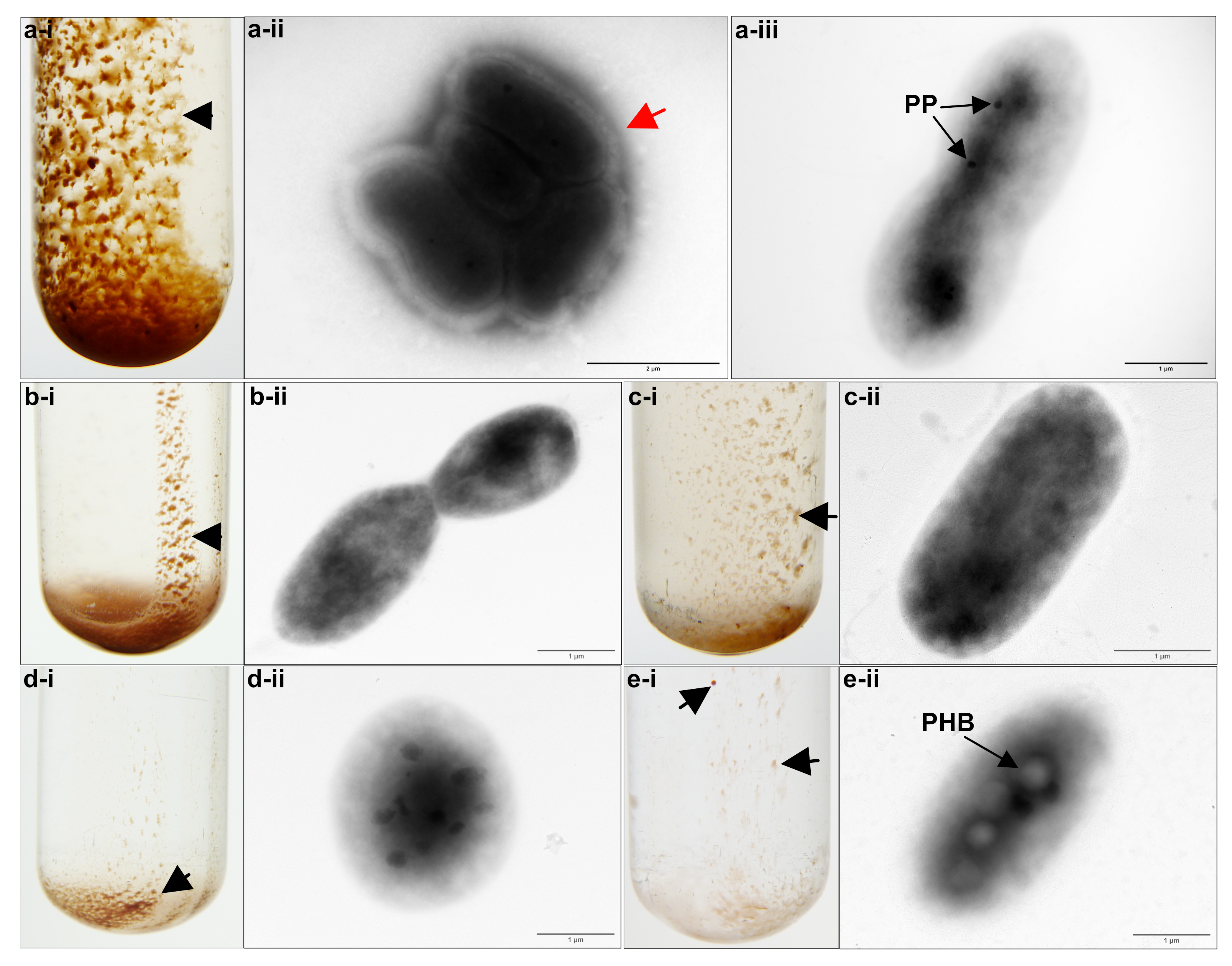

### Supplementary Figure S3

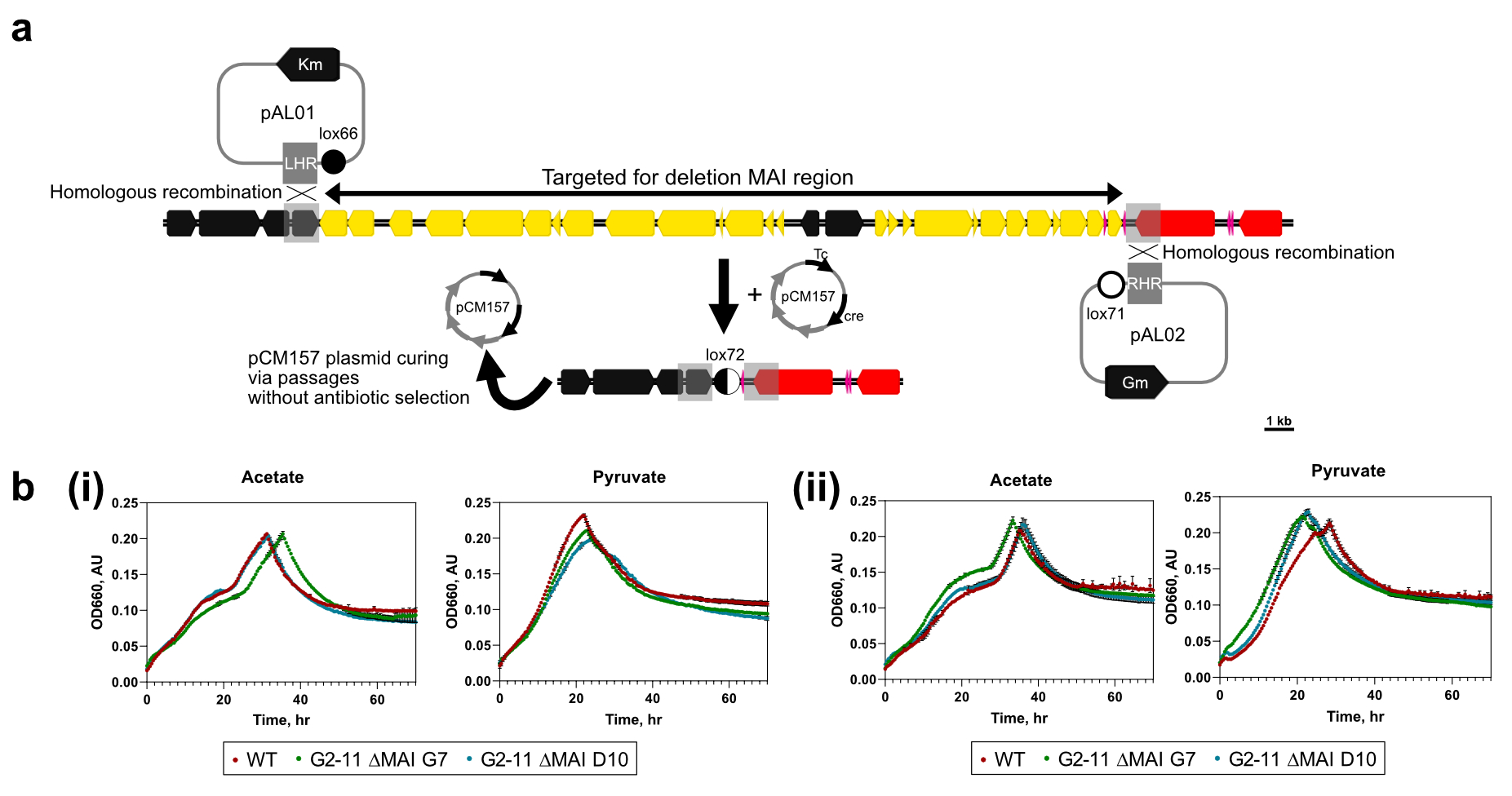

### Supplementary Figure S4

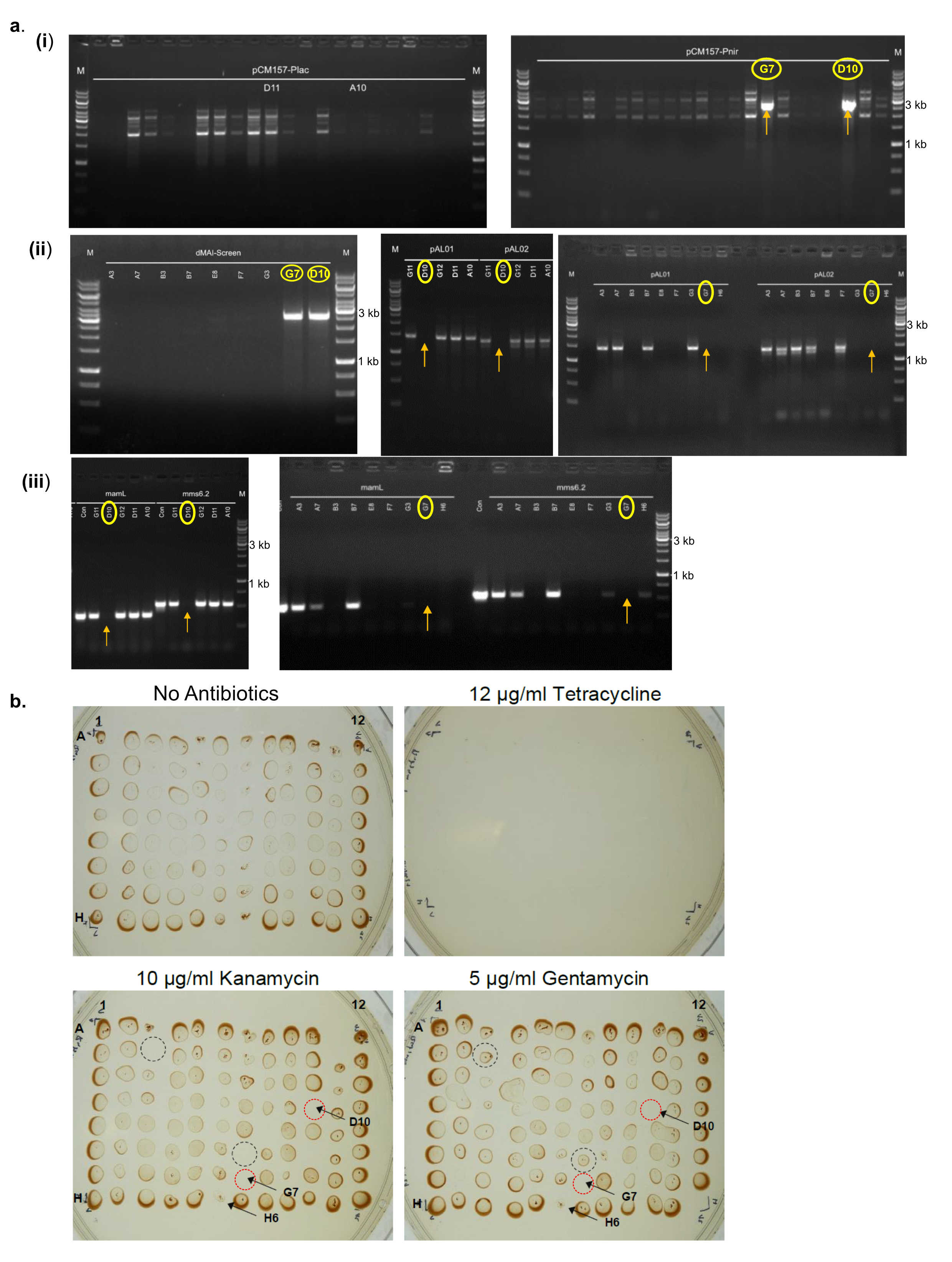

### Supplementary Figure S5

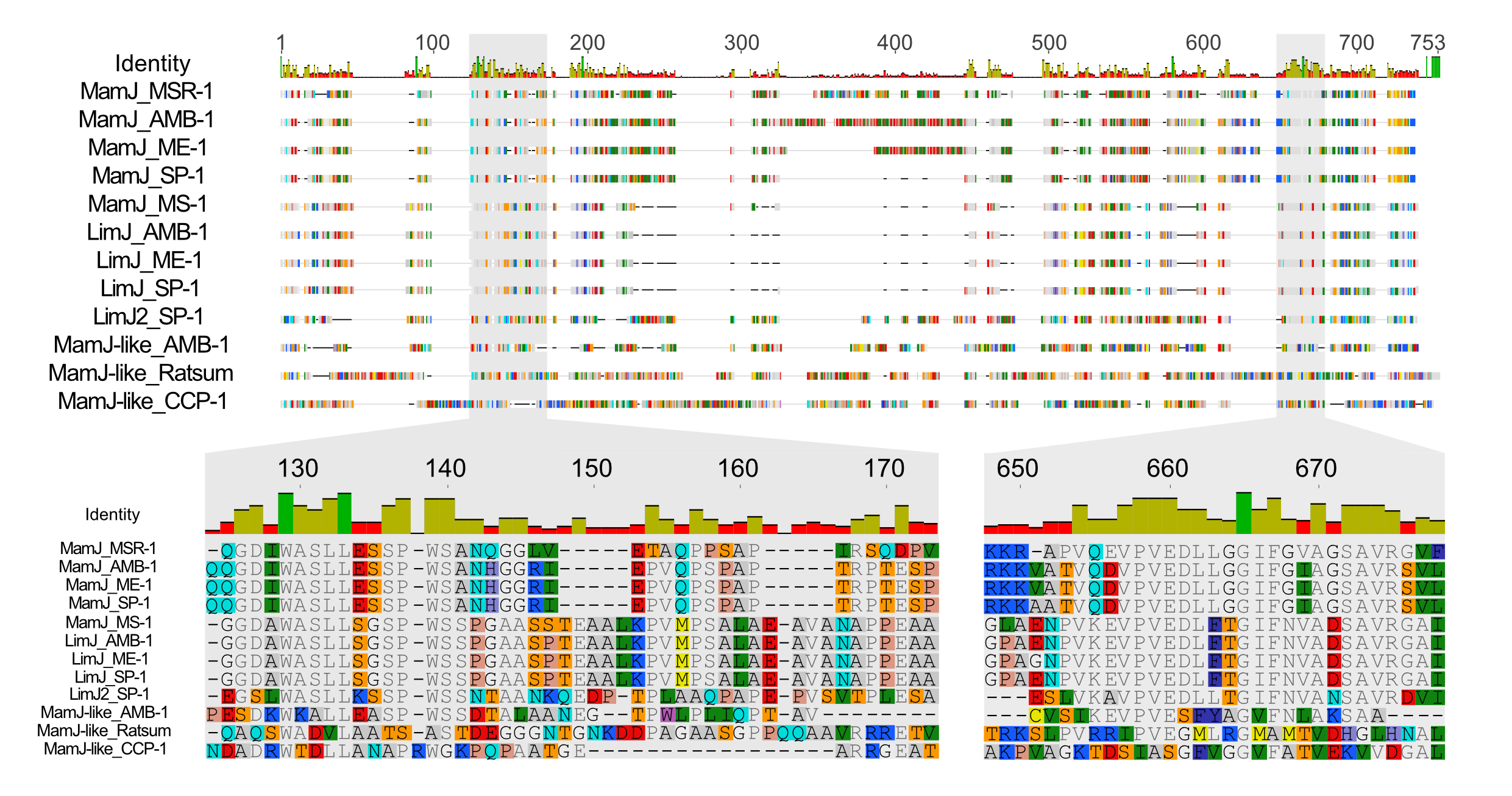

### Supplementary Figure S6

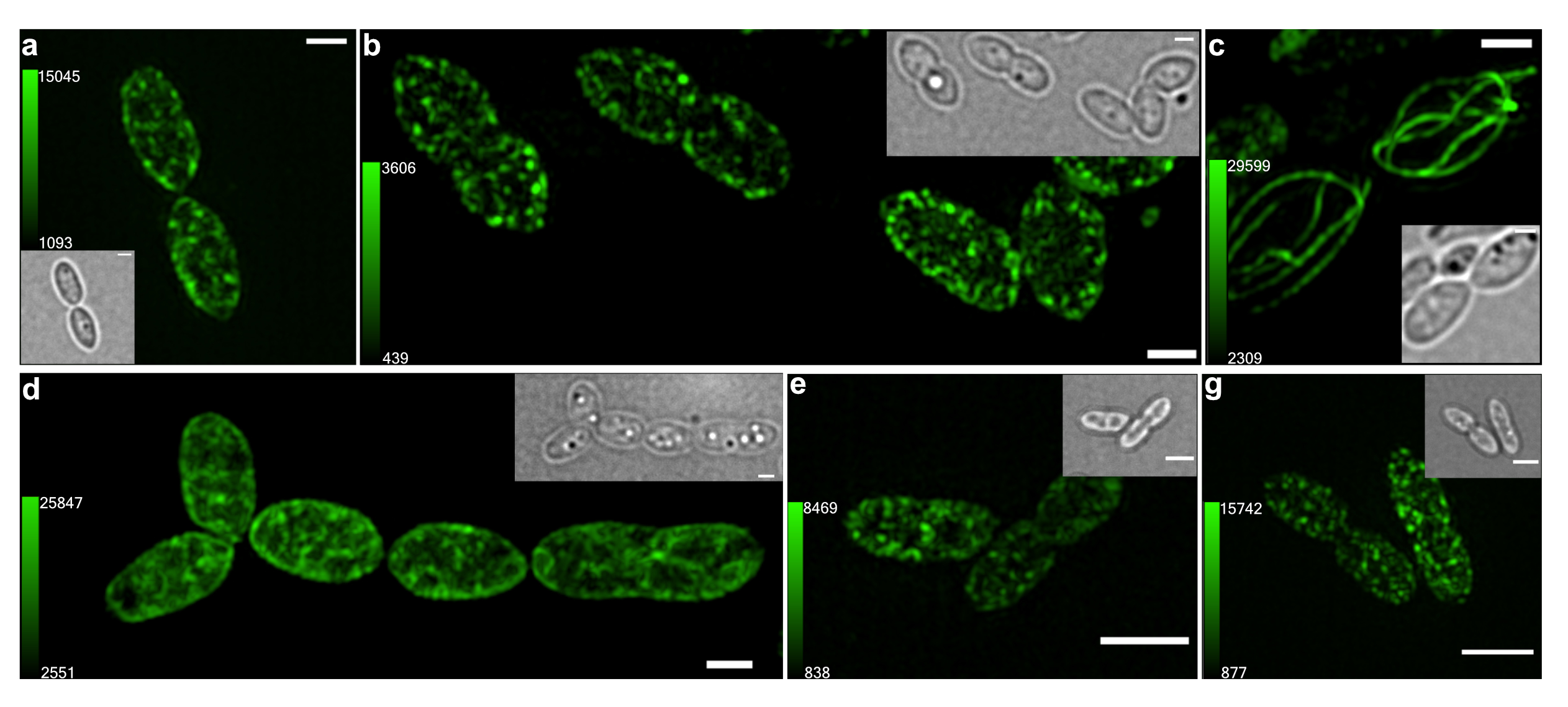
