## Supplementary Table S1 for "Hidden Talents: Silent Gene Clusters Encoding Magnetic Organelle Biosynthesis in a Non-Magnetotactic Phototrophic Bacterium"

**Supplementary Table S1 Results of the BLAST search of the magnetosome proteins encoded in the G2-11 genome against the NCBI database**

| **Gene locus tag** | **Product annotation** | **Best blast hit in cultivated MTB** | **Identity %** | **E value** |
| --- | --- | --- | --- | --- |
| RHOA_v2_0071 | Protein kinase family protein, Mag3 | [protein kinase family protein [Magnetospirillum moscoviense]](https://www.ncbi.nlm.nih.gov/protein/WP_068503441.1?report=genbank&log$=protalign&blast_rank=3&RID=07K8TNWX016), Sequence ID: WP_068503441.1 | 57.32% | 1e-125 |
| RHOA_v2_0072 | 2C domain protein phosphatase, Mag2 | [hypothetical protein [Magnetospirillum moscoviense]](https://www.ncbi.nlm.nih.gov/protein/WP_068503444.1?report=genbank&log$=protalign&blast_rank=7&RID=07KZHYR601R), Sequence ID: WP_068503444.1 | 52.51% | 4e-92 |
| RHOA_v2_0073 | VWA domain-containing protein, Mag1 | [conserved hypothetical protein [Candidatus Terasakiella magnetica]](https://www.ncbi.nlm.nih.gov/protein/CAA7622362.1?report=genbank&log$=protalign&blast_rank=5&RID=07MF0DJB013), Sequence ID: CAA7622362.1 | 68.66% | 4e-134 |
| RHOA_v2_0074 | Magnetosome protein MamH2 | [MFS transporter [Magnetospira sp. QH-2]](https://www.ncbi.nlm.nih.gov/protein/WP_046020676.1?report=genbank&log$=protalign&blast_rank=7&RID=07NAAXEB016), Sequence ID: WP_046020676.1 | 49.30% | 1e-127 |
| RHOA_v2_0075 | Putative magnetosome membrane transporter protein MamO | [magnetosome protein MamO [Rhodospirillaceae bacterium LM-1]](https://www.ncbi.nlm.nih.gov/protein/CAA6606535.1?report=genbank&log$=protalign&blast_rank=9&RID=07NWA3HV013)  Sequence ID: CAA6606535.1 | 47.27% | 1e-160 |
| RHOA_v2_0076 | Magnetosome protein MamM | [magnetosome biogenesis CDF transporter MamM [Magnetospira sp. QH-2]](https://www.ncbi.nlm.nih.gov/protein/WP_046020680.1?report=genbank&log$=protalign&blast_rank=3&RID=07PBH6J4016)  Sequence ID: [WP_046020680.1](https://www.ncbi.nlm.nih.gov/protein/WP_046020680.1?report=genbank&log$=protalign&blast_rank=3&RID=07PBH6J4016) | 49.84% | 1e-111 |
| RHOA_v2_0077 | Magnetosome protein MamL | [hypothetical protein D5085_13130 [Ectothiorhodospiraceae bacterium BW-2]](https://www.ncbi.nlm.nih.gov/protein/QEP43978.1?report=genbank&log$=protalign&blast_rank=5&RID=07PUMTGB013)  Sequence ID: QEP43978.1 | 33.33% | 0.003 |
| RHOA_v2_0078 | Actin-like protein MamK | [actin-like protein MamK [Magnetospira sp. QH-2]](https://www.ncbi.nlm.nih.gov/protein/CCQ72992.1?report=genbank&log$=protalign&blast_rank=5&RID=07RF5PWS01R)  Sequence ID: CCQ72992.1 | 53.87% | 2e-127 |
| RHOA_v2_0079 | Hypothetical protein | No hits found | - | - |
| RHOA_v2_0081 | Magnetosome formation protease MamE | [trypsin-like peptidase domain-containing protein [Magnetovibrio blakemorei]](https://www.ncbi.nlm.nih.gov/protein/WP_069957852.1?report=genbank&log$=protalign&blast_rank=6&RID=07S5N6TC016)  Sequence ID: WP_069957852.1 | 39.65% | 4e-140 |
| RHOA_v2_0082 | Magnetosome protein MamI | [hypothetical protein [Candidatus Terasakiella magnetica]](https://www.ncbi.nlm.nih.gov/protein/WP_069186783.1?report=genbank&log$=protalign&blast_rank=9&RID=07SPPFGS016)  Sequence ID: WP_069186783.1 | 55.36% | 6e-14 |
| RHOA_v2_0083 | Magnetosome protein MamH1 | [MFS transporter [Candidatus Terasakiella magnetica]](https://www.ncbi.nlm.nih.gov/protein/WP_069186784.1?report=genbank&log$=protalign&blast_rank=7&RID=07SXVHHU016)  Sequence ID: WP_069186784.1 | 56.00% | 1e-156 |
| RHOA_v2_0084 | Magnetosome protein MmsF-like1 | [hypothetical protein [Candidatus Terasakiella magnetica]](https://www.ncbi.nlm.nih.gov/protein/WP_069186785.1?report=genbank&log$=protalign&blast_rank=4&RID=07T3KYN5016)  Sequence ID: [WP_069186785.1](https://www.ncbi.nlm.nih.gov/protein/WP_069186785.1?report=genbank&log$=protalign&blast_rank=4&RID=07T3KYN5016) | 57.94% | 1e-44 |
| RHOA_v2_0085 | Magnetosome protein Mms6-like1 | No hits found | - | - |
| RHOA_v2_0090 | Magnetosome protein Mms6-like2 | No hits found | - | - |
| RHOA_v2_0091 | Magnetosome protein MmsF-like2 | [hypothetical protein [Candidatus Terasakiella magnetica]](https://www.ncbi.nlm.nih.gov/protein/WP_069186785.1?report=genbank&log$=protalign&blast_rank=4&RID=07T3KYN5016)  Sequence ID: [WP_069186785.1](https://www.ncbi.nlm.nih.gov/protein/WP_069186785.1?report=genbank&log$=protalign&blast_rank=4&RID=07T3KYN5016) | 47.12% | 2e-28 |
| RHOA_v2_0092 | Iron transporter protein FeoAm | [Fe2+ transport system protein A [Rhodospirillaceae bacterium LM-1]](https://www.ncbi.nlm.nih.gov/protein/CAA6606525.1?report=genbank&log$=protalign&blast_rank=5&RID=07UGWSC2013)  Sequence ID: CAA6606525.1 | 54.79% | 2e-17 |
| RHOA_v2_0093 | Iron transporter protein FeoBm | [Fe2+ transport system protein B [Magnetovibrio blakemorei]](https://www.ncbi.nlm.nih.gov/protein/CAV30756.1?report=genbank&log$=protalign&blast_rank=5&RID=07UPBJCX016)  Sequence ID: [CAV30756.1](https://www.ncbi.nlm.nih.gov/protein/CAV30756.1?report=genbank&log$=protalign&blast_rank=5&RID=07UPBJCX016) | 50.85% | 0.0 |
| RHOA_v2_0094 | Hypothetical protein | No hits found | - | - |
| RHOA_v2_0095 | Magnetosome protein MamP | [magnetochrome domain-containing protein [Magnetovibrio blakemorei]](https://www.ncbi.nlm.nih.gov/protein/WP_069957858.1?report=genbank&log$=protalign&blast_rank=4&RID=07V8CDF5016)  Sequence ID: WP_069957858.1 | 55.77% | 1e-74 |
| RHOA_v2_0096 | Magnetosome protein MamA | [magnetosome protein [alpha proteobacterium LM-1]](https://www.ncbi.nlm.nih.gov/protein/AEX00088.1?report=genbank&log$=protalign&blast_rank=9&RID=07VHZEUV016)  Sequence ID: AEX00088.1 | 42.36% | 3e-50 |
| RHOA_v2_0097 | Magnetosome protein MamQ | [LemA family protein [Magnetospira sp. QH-2]](https://www.ncbi.nlm.nih.gov/protein/WP_046020682.1?report=genbank&log$=protalign&blast_rank=6&RID=07VTPZR0013)  Sequence ID: WP_046020682.1 | 42.86% | 1e-68 |
| RHOA_v2_0098 | Magnetosome protein MamR | [helix-turn-helix domain-containing protein [Magnetospirillum moscoviense]](https://www.ncbi.nlm.nih.gov/protein/WP_068504745.1?report=genbank&log$=protalign&blast_rank=10&RID=07VZWEX8013)  Sequence ID: WP_068504745.1 | 45.24% | 1e-16 |
| RHOA_v2_0099 | Magnetosome protein MamB | [magnetosome biogenesis CDF transporter MamB [Magnetospira sp. QH-2]](https://www.ncbi.nlm.nih.gov/protein/WP_046020684.1?report=genbank&log$=protalign&blast_rank=4&RID=07WDAU27016)  Sequence ID: WP_046020684.1 | 57.64% | 3e-120 |
| RHOA_v2_0100 | Magnetosome protein MamS | [hypothetical protein [Magnetovibrio blakemorei]](https://www.ncbi.nlm.nih.gov/protein/WP_069957862.1?report=genbank&log$=protalign&blast_rank=7&RID=07WHD6Y8016)  Sequence ID: WP_069957862.1 | 46.46% | 6e-32 |
| RHOA_v2_0101 | Magnetosome protein MamT | [magnetochrome domain-containing protein [Magnetospirillum kuznetsovii]](https://www.ncbi.nlm.nih.gov/protein/WP_112147413.1?report=genbank&log$=protalign&blast_rank=5&RID=07WSV2M5016)  Sequence ID: WP_112147413.1 | 61.65% | 4e-48 |
