## Supplementary Table S2 for "Hidden Talents: Silent Gene Clusters Encoding Magnetic Organelle Biosynthesis in a Non-Magnetotactic Phototrophic Bacterium"

**Table S2** Strains used in this study

| **Strain** | **Characteristics** | **Source** |
| --- | --- | --- |
| ***Rhodovastum atsumiense* G2-11** | Wildtype | DSM 21279^1^ |
| ***Rhodovastum atsumiense* ΔMAI** | The region comprising all magnetosome genes, from position 79,446 to 106,923 (27.5 kb) in the chromosome is deleted | This work |
| ***Magnetospirillum gryphiswaldense* MSR-1** | Wildtype, archetype | DSM 6361^2^ |
| ***Magnetospirillum gryphiswaldense* MSR Δ*mamB*** | Δ*mamB* | Lab collection^3^ |
| ***Magnetospirillum gryphiswaldense* MSR Δ*mamM*** | Δ*mamM* | Lab collection^3^ |
| ***Magnetospirillum gryphiswaldense* MSR Δ*mamJ*** | Δ*mamJ* | Lab collection^4^ |
| ***Magnetospirillum gryphiswaldense* MSR Δ*mamK*** | Δ*mamK* | Lab collection^5^ |
| ***Magnetospirillum gryphiswaldense* MSR Δ*mamKY*** | Δ*mamK* Δ*mamY* | Lab collection^6^ |
| ***Magnetospirillum gryphiswaldense* MSR Δ*mamQ*** | Δ*mamQ* | Awal R.P., manuscript in preparation |
| ***Magnetospirillum gryphiswaldense* MSR Δ*mamO*** | Δ*mamO* | Awal R.P., manuscript in preparation |
| ***Magnetospirillum gryphiswaldense* MSR Δ*mamE*** | Δ*mamE* | Awal R.P., manuscript in preparation |
| ***Magnetospirillum gryphiswaldense* MSR Δ*mamI*** | Δ*mamI* | Awal R.P., manuscript in preparation |
| ***Magnetospirillum gryphiswaldense* MSR Δ*mamL*** | Δ*mamL* | Awal R.P., manuscript in preparation |
| ***Magnetospirillum gryphiswaldense* MSR ΔF3** | Δ*mamF* Δ*mmsF* Δ*mmxF* | Uebe R., manuscript in preparation |
| ***E. coli* WM3064** | *thrB1004 pro thi rpsL hsdS* *lacZ*ΔM15 RP4-1360Δ (*araBAD*) 567Δ*dapA1341::[erm pir].* Donor strain for conjugation, auxotroph by DL-α,ε-diaminopimelic acid (DAP) | William Metcalf, UIUC, unpublished |
