## Supplementary Table S3 for "Hidden Talents: Silent Gene Clusters Encoding Magnetic Organelle Biosynthesis in a Non-Magnetotactic Phototrophic Bacterium"

**Table S3 Oligonucleotides used in this study. The sites recognized by restriction enzymes are underlined.**

| **Primer name** | **Sequence 5’-3’** | **Purpose** | **Restriction Enzyme** |
| --- | --- | --- | --- |
| **UNS1-PmamDC fw** | CATTACTCGCATCCATTCTCAGGCTGTCTCGTCTCGTCTCCTTTTTCGCTTTACTAGCTCTTAG | Fluorescent labelling of the selected magnetosome proteins from G2-11 with mNG. The constructs were assembled by Gibson method. | - |
| **mNG100-GSA rev** | GAATTCGCCAGAACCAGCAGCGGAACCTGCTGATCCCTTATACAGTTCGTCCAT |  | - |
| **GSA-MamK(Rat) fw** | GGATCAGCAGGTTCCGCTGCTGGTTCTGGCGAATTCATGAGTGATGAACAGACCGACATT |  | - |
| **UNSX-MamK rev** | GGTGGAAGGGCTCGGAGTTGTGGTAATCTATGTATCCTGGTCAGTCTTTGCCGACGACATC |  | - |
| **PmamDC-mamB(Rat) rev** | CCCACAGGCCCTGCACTGGTTGCTTGTCATATGCTGATCTCCTAAGCTTCG |  | - |
| **PmamDC-mamB(Rat) rev** | CCCACAGGCCCTGCACTGGTTGCTTGTCATATGCTGATCTCCTAAGCTTCG |  | - |
| **PmamDC-mamB(Rat) fw** | CGAAGCTTAGGAGATCAGCATATGACAAGCAACCAGTGCAGGGCCTGTGGG |  | - |
| **GSA-mamB(Rat) rev** | GAATTCGCCAGAACCAGCAGCGGAACCTGCTGATCCTGGCACGCTCCCCGGCAGGGCCTT |  | - |
| **GSA-mamQ(Rat) fw** | GGATCAGCAGGTTCCGCTGCTGGTTCTGGCGAATTCATGGCAAAACCTCCCCCGGA |  | - |
| **UNSX-mamQ(Rat) rev** | GGTGGAAGGGCTCGGAGTTGTGGTAATCTATGTATCCTGGTCATTTGCGATCCCCCATGG |  | - |
| **MamJ-like_Rhatsum_rev** | TATA**GGTACC**TGTCGGCCTCCCAGGCGG | Fluorescent labelling of MamJ-like[G2-11] with GFP | *KpnI* |
| **MamJ-like_Rhatsum_for** | GTAC**CATATG**GCAAAACGGAAAAGGGCCAGG |  | *NdeI* |
| **MmsF-like1_KpnI_for** | ATTC**GGTACC**ATGGATTCCGCTTCGATCGG | Cloning of *mmsF-like1*[G2-11] | *KpnI* |
| **MmsF-like1_SacI_rev** | GTAC**GAGCTC**TCACAGCCGCGCCGCGAGAC |  | *SacI* |
| **MmsF-like2_KpnI_for** | ATTC**GGTACC**ATGATCCCGGGTGAGGAGG | Cloning of *mmsF-like2*[G2-11] | *KpnI* |
| **MmsF-like2_SacI_rev** | GTAC**GAGCTC**TCACAACTTCGCAGCCAGAT |  | *SacI* |
| **RPA1202** | CCCAATTC**CATATG**CCAAGTATCATGATTGGATTGCTCGC | Cloning of *mamI*[G2-11] | *NdeI* |
| **RPA1203** | AAT**CTCGAG**TCAGAGCGATTCGTGGCGCGC |  | *XhoI* |
| **RPA1204** | CCCAATTC**CATATG**CCGATGCGGGTAATCTGGATCCTC | Cloning of *mamL*[G2-11] | *NdeI* |
| **RPA1205** | CGC**CTCGAG**TCATGGGCCAAAATCCACGTCGC |  | *XhoI* |
| **RPA1206** | CCCAATTC**CATATG**GCAAAACCTCCCCCGGACC | Cloning of *mamQ*[G2-11] | *NdeI* |
| **RPA1207** | AAT**CTCGAG**TCATTTGCGATCCCCCATGGTGCG |  | *XhoI* |
| **RPA1208** | GCCAATTC**CATATG**ACAAGCAACCAGTGCAGGGCC | Cloning of *mamB*[G2-11] | *NdeI* |
| **RPA1209** | AAT**CTCGAG**TCATGGCACGCTCCCCGGCAGG |  | *XhoI* |
| **RPA1210** | CCCAATTC**CATATG**AGATACCGGAATTGCCTGACGTGCTCG | Cloning of *mamM*[G2-11] | *NdeI* |
| **RPA1211** | GGC**CTCGAG**TCATCCCTCGCCGGCGACGG |  | *XhoI* |
| **RPA1212** | CCCAATTC**CATATG**GCACCCGACGAGACCGATATCG | Cloning of *mamE*[G2-11] | *NdeI* |
| **RPA1213** | CCG**CTCGAG**TTATGGCATTACAACGAAAAACTCCC |  | *XhoI* |
| **RPA1214** | CCCAATTC**CATATG**GCGGTCCGGCCTTGCCCGG | Cloning of *mamO*[G2-11] | *NdeI* |
| **RPA1215** | GTA**CTCGAG**TCAGTGGCCGCCGATGAACATC |  | *XhoI* |
| **RPA1254** | GGGAATTC**CATATG**AGTGATGAACAGACCGACATTG | Cloning of *mamK*[G2-11] | *Ndel* |
| **RPA1255** | CTTTTCGAGCACATCGTCGCCAAC |  |  |
| **RPA1256** | GTGCTCGAAAAGCAATCGTTTCTGGAATTG |  |  |
| **RPA1257** | CAT**CTCGAG**TCAGTCTTTGCCGACGACATC |  | *XhoI* |
| **KpnI-dMAI(Ratsu)-LHR fw** | GAA**GGTACC**GTTCGCCGGCATCCGCCA | Deletion of the MAI | *KpnI* |
| **NotI-dMAI(Ratsu)-LHR rev** | TTT**GCGGCCGC**CGGTCAGGGCGCAGGGTC |  | *NotI* |
| **BamHI-dMAI(Ratsu)-RHR fw** | GAA**GGATCC**CAGGGATGGGCGCAAACG |  | *BamHI* |
| **KpnI-dMAI(Ratsu)-RHR rev** | CTT**GGTACC**TGAAACCGGGGCGCCAGC |  | *KpnI* |
| **dMAI(Rat) fw** | GGGAATATCTCATGGGCTTCGATC |  |  |
| **dMAI(Rat) rev** | GGAGAGCACTGTGTAGGATAGG |  |  |
| **pAL01 rev** | CAGGAAACAGCTATGACCATGATTAC |  | - |
| **pAL02 fw** | CCTGCATATCCCGATTCAACGGC |  | - |
