## Supplementary Table S4 for "Hidden Talents: Silent Gene Clusters Encoding Magnetic Organelle Biosynthesis in a Non-Magnetotactic Phototrophic Bacterium"

**Table S4 Plasmids used in this study**

| **Plasmid** | **Characteristics** | **Source** |
| --- | --- | --- |
| **pBamII-Tn5-Tc** | *TcR, AmpR*, *p15A ori*, *mini-Tn5*; general vector used for expression of the individual mNG-tagged proteins. Suicide vector, a cassette is introduced by random chromosomal insertion mediated by mini-Tn5 | Uebe R., manuscript in preparation |
| **pBamII-Tn7-Km** | *KmR, AmpR, p15A ori, Tn7, tr2*, *T1*, general vector used for expression of the individual untagged genes. Suicide vector, a cassette is introduced by chromosomal insertion mediated by Tn7 into the *attTn7* site. | Uebe R., manuscript in preparation |
| **pBamII-Tn5-mNG-mamK** | pBamII-Tn5-Tc with inserted P*mamDC*-*mNG-mamK[G2-11]* | This work |
| **pBamII-Tn5-mNG-mamQ** | pBamII-Tn5-Tc with inserted P*mamDC*-*mNG-mamQ[G2-11]* | This work |
| **pBamII-Tn5-mamK-mNG** | pBamII-Tn5-Tc with inserted P*mamDC*-*mamB[G2-11]-mNG* | This work |
| **pBamII-Tn7-mamI** | pBamII-Tn7-Km with inserted P*mamDC*-*mamI[G2-11]* | This work |
| **pBamII-Tn7-mamL** | pBamII-Tn7-Km with inserted P*mamDC*-*mamL[G2-11]* | This work |
| **pBamII-Tn7-mamQ** | pBamII-Tn7-Km with inserted P*mamDC*-*mamQ[G2-11]* | This work |
| **pBamII-Tn7-mamB** | pBamII-Tn7-Km with inserted P*mamDC*-*mamB[G2-11]* | This work |
| **pBamII-Tn7-mamO** | pBamII-Tn7-Km with inserted P*mamDC*-*mamO[G2-11]* | This work |
| **pBamII-Tn7-mamM** | pBamII-Tn7-Km with inserted P*mamDC*-*mamM[G2-11]* | This work |
| **pBamII-Tn7-mamE** | pBamII-Tn7-Km with inserted P*mamDC*-*mamE[G2-11]* | This work |
| **pAL01-MCS1-KmR** | *KmR, gusA, lacZ, lox71,* MCS from pBBR-MCS5*, pK19mobGII* vector backbone | ^1^ |
| **pAL02/2-MCS2-GmR** | *GmR, lox66,* MCS from pBBR-MCS5*, pT18mob2* vector backbone | ^1^ |
| **pAL01-LHR** | pAL01-MCS1-KmR with inserted left homology region of the MAI in G2-11 | This work |
| **pAL02-RHR** | pAL02/2-MCS2-GmR with inserted right homology region of the MAI in G2-11 | This work |
| **pCM157-P*_lac_*** | *TcR,* expression of Cre-recombinase under control of P*lac* | ^2^ |
| **pCM157-P*_nir_*** | *TcR,* expression of Cre-recombinase under control of P*nir* | ^3^ |
| **pTpsMAG1** | *KmR, CmR*, *p15A ori*, mariner *tps* *mamAB*, *mamGFDC*, *mms6*, *mamYXZftsZm*, *feoAB1* | ^4^ |
